# Neural stem cells alter nucleocytoplasmic partitioning and accumulate nuclear polyadenylated transcripts during quiescence

**DOI:** 10.1101/2021.01.06.425462

**Authors:** A. Rossi, A. Coum, M. Madelenat, L. Harris, A. Miedzik, S. Strohbuecker, A. Chai, H. Fiaz, R. Chaouni, P. Faull, W. Grey, D. Bonnet, F. Hamid, E. V. Makeyev, A. P. Snijders, G. Kelly, F. Guillemot, R. Sousa-Nunes

## Abstract

Quiescence is a cellular state characterised by reversible cell-cycle arrest and diminished biosynthetic activity that protects against environmental insults, replicative exhaustion and proliferation-induced mutations^1^. Entry into and exit from this state controls development, maintenance and repair of tissues plus, in the adult central nervous system, generation of new neurons and thus cognition and mood^2–4^. Cancer stem cells too can undergo quiescence, which confers them resistance to current therapies^5, 6^. Despite clinical relevance, quiescence is poorly understood and is defined functionally given lack of molecular markers. Decrease of the most resource-intensive cellular process of protein synthesis is a feature of quiescence, controlled across species and cell types by inhibition of the Target of Rapamycin (TOR) pathway^1, 7^. Here, we combine *Drosophila* genetics and a mammalian model to show that altered nucleocytoplasmic partitioning and nuclear accumulation of polyadenylated RNAs are novel evolutionarily conserved hallmarks of quiescence regulation. Furthermore, nuclear accumulation of messenger RNA (mRNA) in quiescent NSCs (qNSCs) largely predicts protein downregulation, accounting for uncoupling between transcriptome and proteome in quiescence. These mechanisms provide a previously unappreciated regulatory layer to reducing protein synthesis in quiescent cells, whilst priming them for reactivation in response to appropriate cues.

---

NSCs give rise to various neuronal and glial cell types in the central nervous system (CNS), mostly during development but also during adult stages in many species, including humans^8–12^. In adult mammals, NSCs are mostly quiescent^13, 14^ and their exit from quiescence (reactivation) is controlled by cell-extrinsic and -intrinsic mechanisms in response to physiological and behavioural stimuli including physical exercise, novel environments, social interactions, and diet^4^. Across cell types, a variety of intercellular signals, receptors and downstream pathways converge on the TOR pathway for quiescence regulation in eukaryotic cells^1^. We and others have shown that in NSCs from fruitflies to mammals, the TOR pathway integrates aminoacid availability and the nutritionally-regulated Insulin / Insulin Growth Factor signalling pathway towards the quiescence/activation decision via downstream effectors such as forkhead-box FoxO transcription factors and ribosomal protein S6 kinase, which control cell-cycle and protein translation^15–20^.

Quiescence is heterogeneous, consisting of a continuum of states between near-active (“shallow”) and profound quiescence (dormancy), with depth defined by reactivation speed^21–23^. A bistable molecular network converts graded cues into ON or OFF S-phase entry^23, 24^. In the fruitfly *Drosophila melanogaster*, the cell-cycle stage at which divisions pause also contributes to heterogeneity of qNSCs, arrested in either G1 or G2^25^.

*Drosophila* NSCs are a powerful model with which to study fundamental molecular and cellular mechanisms of stem cell properties, including quiescence^26^. Out of ∼100 pairs of central brain NSCs plus hundreds in the ventral nerve cord, all but five pairs undergo quiescence during a period of ∼24-48 hours that intervenes between embryonic and postembryonic neurogenesis^27, 28^. In addition to paused proliferation, *Drosophila* qNSCs differ from active NSCs (aNSCs) in a number of discernible ways: size (soma diameter ∼4 μm in early larvae, when quiescent and ∼10-12 μm in late larvae, when active); morphology (presenting a cytoplasmic extension/fibre of unknown function only when quiescent^27^); expression levels of NSC markers (e.g. downregulating the cortical protein Miranda (Mira) and the HES family basic helix-loop-helix transcriptional repressor Deadpan (Dpn) below detectability in a considerable fraction of qNSCs) (Fig. 1a,d). Following larval hatching and feeding, qNSCs reactivate in a stereotypical spatiotemporal pattern^15, 27^. As they reactivate, the qNSC soma enlarges, fibres thicken and are lost (via inheritance by the first post-reactivation transit-amplifying daughter cell^29^), and NSC markers become detectable (Fig. 1d and Extended Data Fig. 1). At late larval stages, all NSCs are active. Together, these properties enable fast and quantitative study of the processes governing NSC quiescence/reactivation in *Drosophila*.

**Fig. 1.**
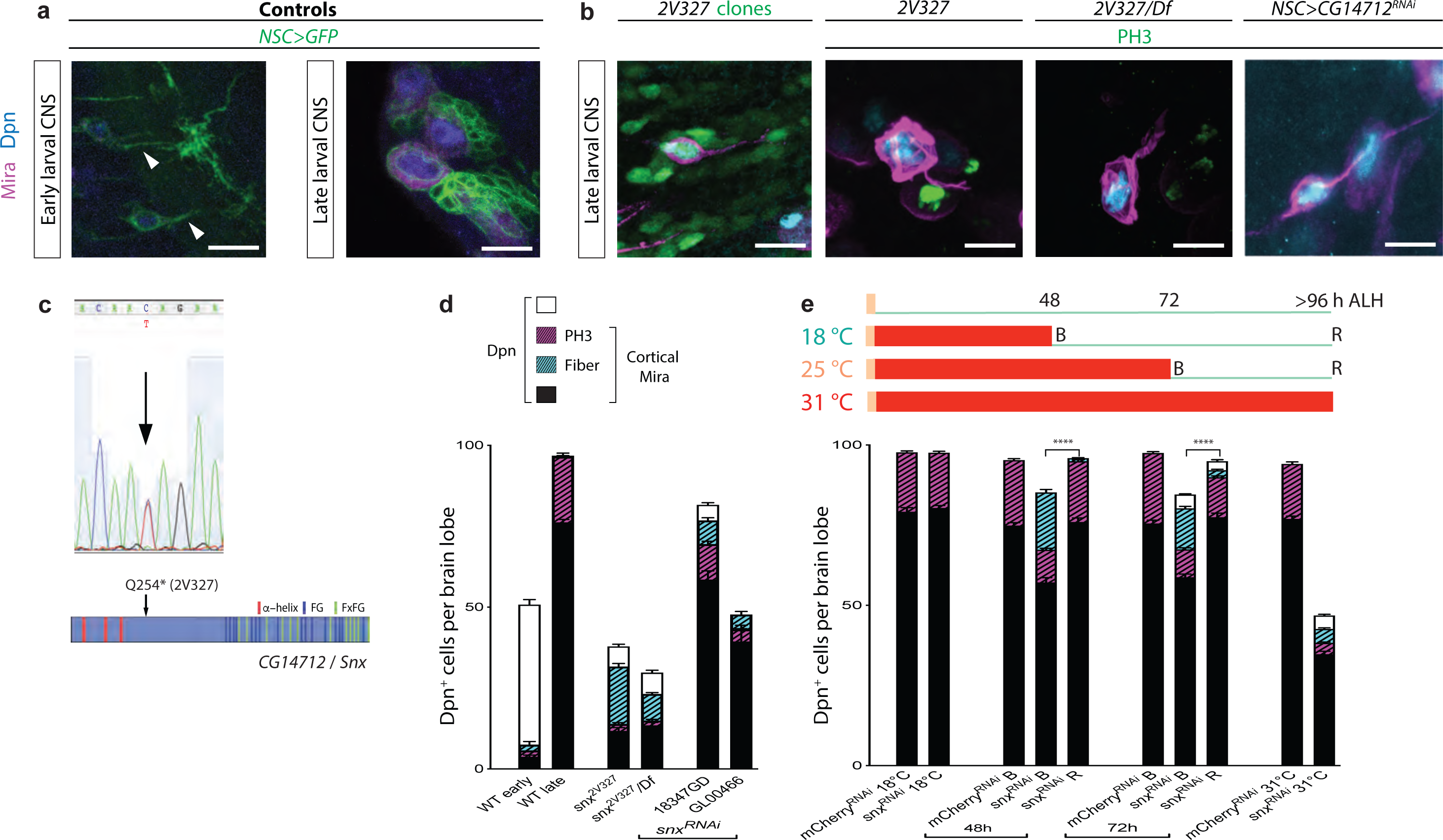
Downregulation of novel *Drosophila* protein CG14712/Snx induces anachronical qNSCs. **a**, WT qNSCs are present in early but not late larval CNSs, display a cytoplasmic fibre (arrowheads) and downregulate Mira and Dpn. **b**, Late larval NSCs of 2V327 mutant or with CG14712 knock-down (shown is RNAi GL00466) have a morphology reminiscent of qNSCs and are negative for the mitotic marker PH3. Images are maximum intensity projections of a few optical sections; scale bars: 10 µm. **c**, 2V327 mutants contain a premature STOP codon at aminoacid position 254 in CG14712/Snx, whose structural motifs are indicated. **d**, Quantification of detectable Dpn^+^ cells per brain lobe of early (newly-hatched) and late (third-instar) WT larvae as well as late *snx* mutant and *NSC>snx^RNAi^* larvae (two independent RNAis). Dpn^+^ cells were further scored for Mira (black bars – full and hatched) and, within these, for expression of PH3 and presence of a fibre (note that with this marker combination fibres are only revealed if cells express detectable cortical Mira). Histograms represent mean and error bars s.e.m. **e**, Schematic of experimental design to transiently induce RNAi expression in larval NSCs via temperature shifts: none/low at 18 °C and maximal at 31 °C. ‘B’ refers to animals scored Before or after Recovery ‘R’ period (recovery meaning animals placed at 18 °C to abolish RNAi induction); ALH, after larval hatching. Quantifications as per d. Student’s t-test was performed to compare number of Dpn^+^ cells in the conditions indicated; ****p<0.0001.

The nuclear pore complex (NPC) is the evolutionarily conserved gateway for bidirectional transport between nucleus and cytoplasm in eukaryotic cells. The NPC is assembled from multiple copies of ∼30 distinct nucleoporins^30^ grouped into several major classes (Supplementary Table 1). Around one third are scaffold nucleoporins, which form two eight-fold rotationally-symmetric doughnut-shaped structures outlining the pore canal^30^. Another third are FG-nucleoporins, containing repeated phenylalanine (F) and glycine (G) motifs. FG and FxFG-rich sequences promote natively unfolded conformations that extend either to the centre of the pore canal in “mesh” or “barrier” nucleoporins; or into the cytoplasm or nucleoplasm in “asymmetric” nucleoporins. FG nucleoporins form highly-specific low-affinity interactions with cargo complexes to be transported, whilst excluding unwanted ones^30, 31^. Nucleoporins can take on diverse functions besides their role in nucleocytoplasmic transport, most notably transcriptional and microtubule regulation^32–34^.

Whilst small molecules (<40 kDa) can passively diffuse across the NPC, efficient distribution of larger proteins between nucleus and cytoplasm depends on active transport. This is fuelled by hydrolysis of guanosine nucleoside triphosphate (GTP) by the small GTPase Ran^35^. In this case, evolutionarily conserved karyopherins (called importins or exportins, Supplementary Table 2) associate with nuclear localisation and/or nuclear export signals in protein cargo to facilitate their translocation across the NPC^36^.

Most mRNAs are exported from the nucleus to the cytoplasm as messenger ribonucleoprotein complexes (mRNPs) in a manner reliant on ATP-dependent events rather than Ran GTPase and karyopherins^37^. Following pre-messenger RNA processing, which typically includes 5’-capping, splicing, 3’-cleavage and polyadenylation, mature mRNPs associated with components of the TRanscription and Export (TREX), and Nuclear RNA export factor 1 / Nuclear Transport Factor 2 Like Export Factor 1 (NXF1/NXT1) complexes are irreversibly translocated to the cytoplasm^37^. In addition to the non-discriminatory bulk mRNA export pathway, metazoans have evolved selective mRNA export pathways, less well characterised. These employ recruitment of RNA binding proteins via post-transcriptional modifications, structural and/or *cis* elements within the mRNA (named untranslated sequence elements for regulation, USER, codes; though some are embedded in coding sequence). USER codes promote coordinated export of functionally related mRNAs yet a single transcript can have multiple USER codes, which can synergise or compete, so some mRNAs may utilise several possible export pathways in accordance with export factor availability^37–39^. Some selective mRNA export mechanisms depend on specific nucleoporins or Exportin-1 (Xpo1/Crm1)^37, 38^.

It is emerging that levels and/or complement of nucleoporin and karyopherin complexes vary between cell and tissue types^40–42^. Here, we expand this to aNSCs versus qNSCs, demonstrating a previously unappreciated layer of gene expression regulation for the transition between active and quiescent states.

## Downregulation of novel *Drosophila* protein induces anachronic qNSCs

In a forward-genetic ethyl methane-sulfonate screen, we recovered a *Drosophila* mutant (2V327) in which late larval NSCs, normally active, were cell-cycle arrested whilst displaying a fibre, features of qNSCs. This was seen in whole homozygous animals and homozygous clones induced in early larvae (Fig. 1b) demonstrating cell-autonomy of the phenotype and derivation from mitotic recombination (basis of labelled clone generation).

We hypothesized that the 2V327 mutation led to anachronic quiescence reentry of NSCs. Deficiency-mapping exposed a small genomic region responsible for the phenotype, with hemizygote animals recapitulating that of homozygotes (Fig. 1b and Extended Data Fig. 2). Amongst seven protein-coding genes within the candidate region, RNA interference (RNAi) for only one, *CG14712*, phenocopied the 2V327 mutant (Fig. 1b). Genomic sequencing of *CG14712* exons in 2V327 heterozygous animals uncovered a premature STOP codon, consistent with its disruption causing the phenotype (Fig. 1c). We named this previously uncharacterized gene *snorlax* (*snx*). Homozygous and hemizygous *snx^2V3^*^27^ animals were both lethal as undersized late larvae and quantification of NSC features showed comparable phenotypes (Fig. 1d). 2V327 is thus a strong loss-of-function allele, likely a null. Appreciably fewer than the customary ∼100 cells per central brain lobe were detectable with Dpn and Mira antibodies in *snx^2V327^* (Fig. 1d) but staining for the apoptotic marker Death Caspase-1 (Dcp-1) was negative in all NSCs examined (Extended Data Fig. 3). Overall, the presence of a fibre accompanying cell-cycle arrest, along with detection of fewer Dpn and Mira-positive cells, is consistent with late larval quiescence in *snx* mutant and knockdown NSCs.

**Fig. 2.**
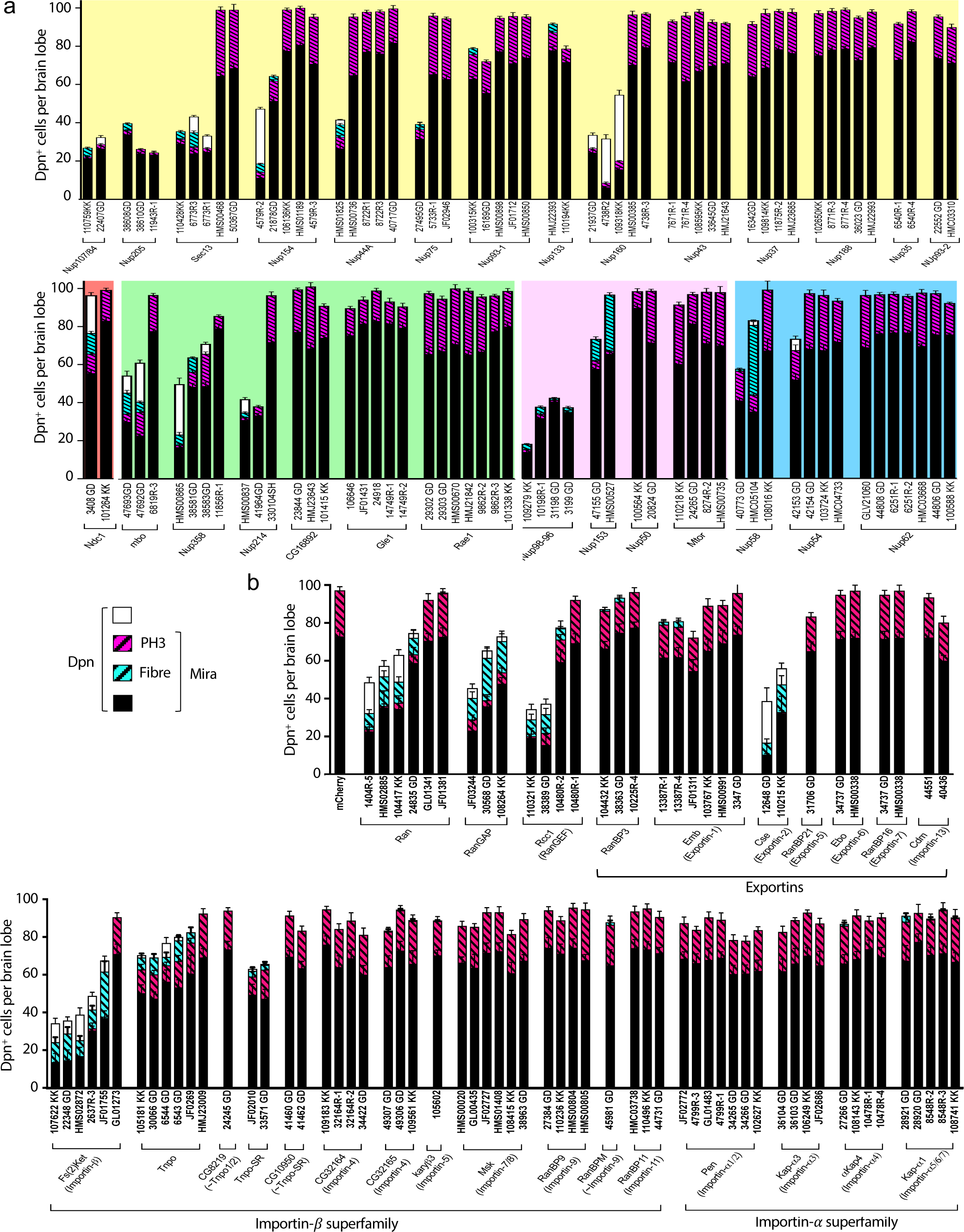
Downregulation of various nucleocytoplasmic transport regulators in *Drosophila* NSCs induces quiescence features anachronically in late larvae. a, Nucleoporins (background colour-coded by structural class − see Extended Data Table 1). **b**, Ran, its guanine nucleotide exchange factors and karyopherins. Wherever possible, multiple RNAis (each identified under their respective histogram) were used against the same target. Quantifications as per Fig. 1d. Histograms represent mean and error bars s.e.m..

**Fig. 3.**
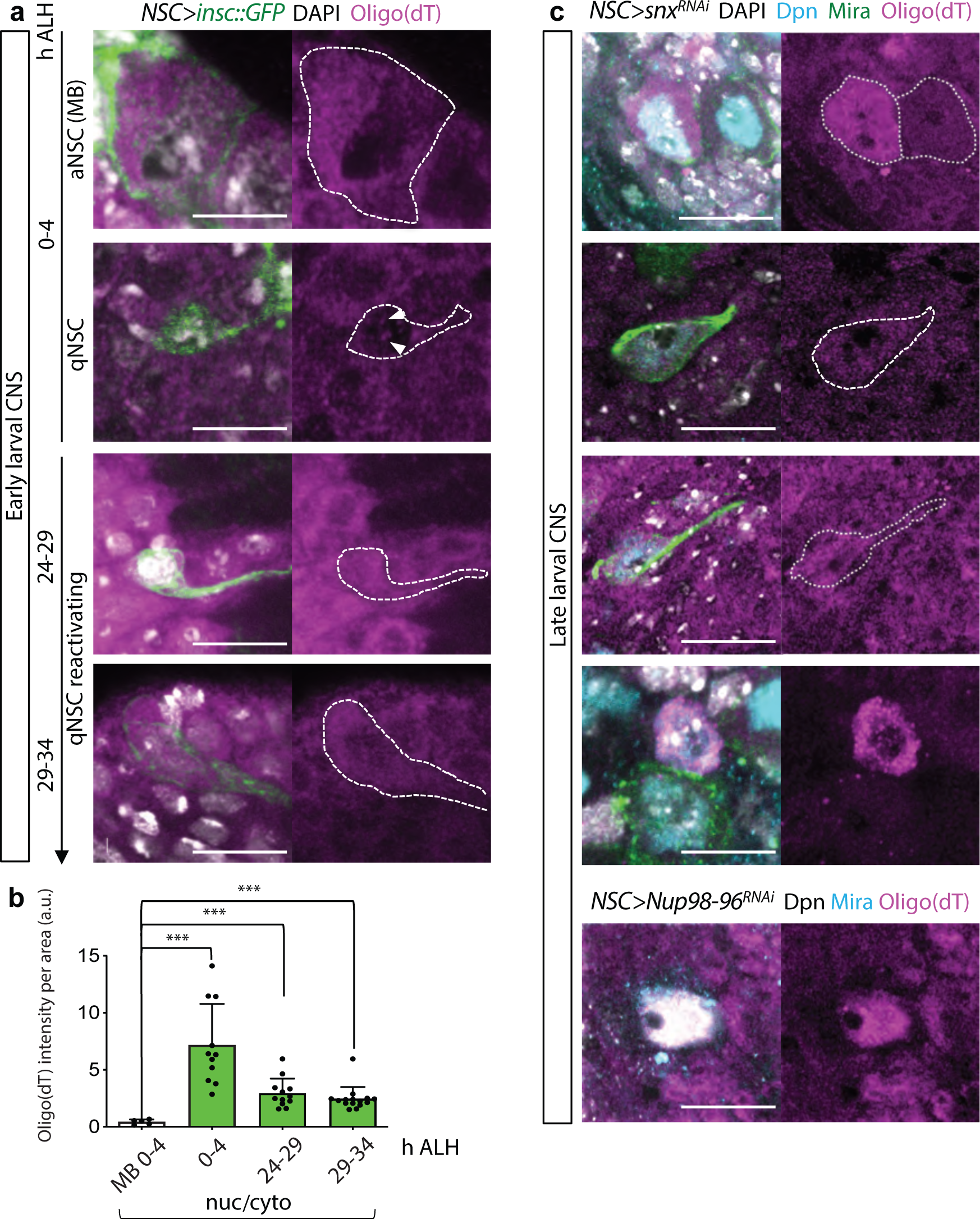
Drosophila qNSCs accumulate nuclear poly(A) RNA relative to aNSCs. **a,** Permanently active NSCs (mushroom body neuroblasts, MB) are larger and contain noticeably more poly(A) RNA than deeply quiescent NSCs (0-4 h after larval hatching, ALH). Following the same qNSC over time (across specimen) (Extended Data Fig. 1), revealed gradual increase of poly(A) RNA as reactivation progressed. Arrowheads point at poly(A) RNA accumulations within the nucleus. **b**, Quantifications from individual cells such as those depicted in a. Histograms represent mean and error bars s.e.m. of values normalised to MB average. Mann-Whitney test, ***p≤0.0005. **c**, Nuclear accumulation of poly(A) RNA seen in some NSCs with Snx or Nup98-96 knockdown, particularly in deeply quiescent NSCs as reported by nuclear Mira. Images are maximum intensity projections of a few optical sections (split magenta channel with hatched outline of NSCs when cell cortex was discernible); scale bars: 10 µm.

Reversibility is a defining feature of quiescence. To test reversibility of the *snx* loss-of-function phenotype in NSCs, we transiently induced *snx^RNAi^* in these cells followed by a period of recovery, and analysed animals before (B) and after recovery (R) (Fig. 1e). Cells undetectable by anti-Mira or anti-Dpn before recovery re-emerged as visible with these markers following the recovery period, demonstrating that they had neither died nor differentiated. Furthermore, recovery also decreased the number of NSCs displaying fibres and increased the mitosis index (reported by expression of phospho-histone H3, PH3), indicating shift of qNSCs to aNSCs (Fig. 1e). We concluded that the NSC phenotype of *snx^RNAi^* was reversible and that, by all criteria, *snx* downregulation in NSC induces anachronic quiescence.

## Specific attenuations of nucleocytoplasmic transport factors in *Drosophila* NSCs induce quiescence features

CG14712/Snx is a putative FG nucleoporin (Fig. 1c; flybase.org), most similar to Nup98, which is found on both the nuclear and cytoplasmic sides of the NPC^30, 43^ (Supplementary Table 1). There is a *bona fide Drosophila* Nup98, however, encoded by the *Nup98-96* gene (Nup98 and Nup96 are generated from a single transcript as a polypeptide precursor that undergoes proteolytic cleavage in various species^44^). We found that like for *snx*, transient knockdown in NSCs of *Nup98-96* followed by recovery also induced quiescence as defined by reversible downregulation of Dpn, Mira and PH3 accompanied by presence of a fibre (Extended Data Fig. 4).

**Fig. 4.**
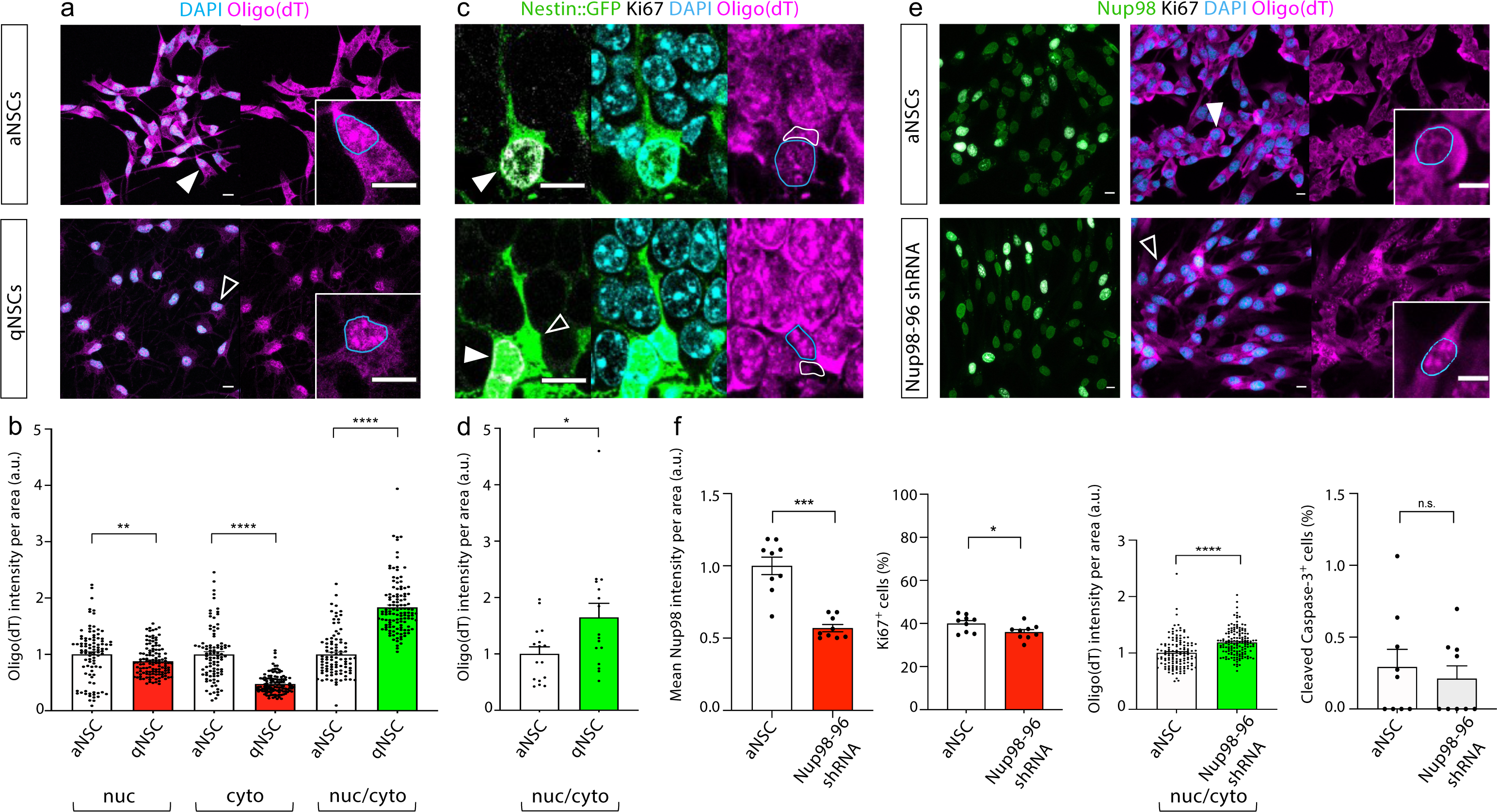
Mouse qNSCs accumulate nuclear poly(A) RNA relative to aNSCs. **a**, **e**, Adult mouse adult hippocampal NSC cultures; quiescence induced by BMP4. Insets in **a** are single-channel high-magnification of single cells (arrowheads in low-magnification) with blue nucleus outline. **b**, Quantifications from individual cells such as those depicted in a. Histograms represent mean and error bars s.e.m. of values normalised to respective aNSC average. Mann-Whitney test, **p<0.005, ****p<0.0001. **c**, Adult mouse hippocampal NSCs *in vivo* identified by Nestin::GFP, with active and quiescent identified by presence/absence of Ki67 (filled and open arrowheads, respectively); nuclear and cytoplasmic domains outlined in blue and white, respectively. **d**, Quantifications from individual cells such as those depicted in c. Histograms represent mean and error bars s.e.m. of values normalised to aNSC average. Mann-Whitney test, *p<0.05. **e**, Nup98 knockdown induces quiescence features such as reduction in Ki67^+^ cells and increase of nucleocytoplasmic ratio of poly(A) RNA. **f**, Quantifications from specimen such as those depicted in e. Histograms represent mean and error bars s.e.m.; first two graphs are values normalised to average in untransfected cells (Nup98 intensity within nuclei). Mann-Whitney test, ****p<0.0001. Images are maximum intensity projections of a few optical sections; scale bars: 10 µm.

We next wondered whether qNSC induction was specific to knockdown of Nup98-like factors or might result also from downregulation of other nucleoporins. We tested available RNAi lines against the other *Drosophila* nucleoporins spread across structural classes. NSC-specific knock-down of 17 out of 27 further nucleoporins induced features of quiescence (fewer Dpn, Mira and PH3-positive cells, plus fibres) with at least one RNAi, 12 of which with more than one (Fig. 2a). Negative RNAi outcomes could be due to the target having no role in quiescence regulation or to maternal contribution^45^, long protein half-life (reported for scaffold nucleoporins^46^) and/or ineffective RNAi (although some constructs that did not induce a phenotype in our assay appeared effective in other contexts^47–49^). Nonetheless, the fact that knockdown of multiple nucleoporins among all classes induced features of quiescence suggested that disruption of their function at the NPC caused the phenotype.

To verify that nucleocytoplasmic transport perturbation underlined NSC quiescence induction, we knocked down active transport components. Knockdown of Ran, its GTPase activating protein (RanGAP) and guanine nucleotide exchange factor (RanGEF), or of a small subset of karyopherins induced qNSC features (Fig. 2b). In particular, knockdown of Exportin-1 and -2, Importin-ß, Tnpo and Tnpo-SR led to strong phenotypes with at least two RNAis (Fig. 2b). Thus, specific perturbations of nucleocytoplasmic transport induce quiescence in *Drosophila* NSCs.

## WT and induced *Drosophila* qNSCs accumulate nuclear polyadenylated RNA

Karyopherins have been shown to mediate transport of functionally-related proteins^36, 50^. Examining biological process gene ontology (GO) terms of known cargo for specific karyopherins^36, 50^ whose knockdown induced qNSCs (Fig. 2b) led us to hypothesise that messenger RNA (mRNA) processing might be altered in quiescent versus active NSCs. Furthermore, we noted that Nups with analogous phenotype (Fig. 2a) included those of the so-called mRNA export platforms^30^. We thus considered whether there might be altered distribution of polyadenylated (poly(A)) RNA between aNSCs and qNSCs.

*In situ* hybridisation with an oligo(dT) probe reported visibly lower poly(A) in qNSCs than aNSCs, as expected from diminished transcription in quiescent cells (Fig. 3a). Nonetheless, we reproducibly found discrete poly(A) accumulations within nuclei of deeply quiescent NSCs (newly-hatched larvae), whereas in permanently active mushroom body NSCs nuclear poly(A) localised predominantly at the nuclear periphery (Fig. 3a). We quantified relative poly(A) immunofluorescence in nuclear and cytoplasmic compartments of NSCs in the two states and discovered that the nuclear/cytoplasmic ratio of poly(A) was higher in qNSCs than in the mushroom body NSCs, which never undergo quiescence (Fig. 3b). In case there was something particular about permanently-active NSCs, we determined the nuclear/cytoplasmic ratio of poly(A) in the same NSC as it reactivated (identifying the same cell across specimen). Therein, we found decrease in relative nuclear accumulation of poly(A) as the NSC reactivated (Fig. 3b), consistent with aNSCs having lower nuclear/cytoplasmic ratio of poly(A) than qNSCs. We concluded that relative accumulation of nuclear poly(A) RNA is a trait of qNSCs in *Drosophila*.

To further assess similarity between physiological quiescence and quiescence induced by attenuation of nucleocytoplasmic transport factors, we examined whether relative accumulation of nuclear poly(A) RNA also occurred in the latter. Consistent with quiescence reentry as opposed to quiescence maintenance, induced anachronic qNSCs were generally larger and more loaded with poly(A) relative to the deeply quiescent NSCs of newly-hatched larvae. Inspection of late larval NSCs with knockdown of either Snx or Nup98-96 revealed cells with very strong nuclear poly(A) signal (Fig. 3c).

## Nuclear accumulation of polyadenylated RNA is a hallmark of quiescence

We reasoned that nuclear accumulation of transcripts would be an efficient way for quiescent cells to reduce protein synthesis whilst remaining able to quickly resume it in response to reactivation cues. To test if this might be a widespread mechanism of quiescence, we first examined primary cultures of adult mouse hippocampal NSCs, where quiescence is induced by addition of BMP4, mimicking a niche signal^51, 52^. We had previously determined that these cells become quiescent after 3 days in BMP4^52^ and used this timepoint as the quiescent condition unless otherwise specified. Synchronicity of quiescence induction in culture permitted determination of relative levels of nuclear and cytoplasmic poly(A) across many cells in an equivalent state. We found that both nuclear and cytoplasmic poly(A) signals decreased in qNSCs compared to aNSCs, but that cytoplasmic levels consistently decreased more, resulting in increased nuclear/cytoplasmic ratio (Fig. 4a,b). Importantly, inspection of adult hippocampal NSCs *in vivo* also showed increased nuclear/cytoplasmic poly(A) ratio in quiescent versus active cells (Fig. 4c,d). Separation between the two conditions was less marked in tissue as there is continuity between the two states and no means of determining how long each cell had been quiescent or active for at the time of fixation. Notwithstanding, the extent of relative poly(A) RNA accumulation was remarkably similar to that found *in vitro*. Surveying of human blood marrow haematopoietic stem and progenitor cells returned the same finding (Extended Data Fig. 5). We concluded that nuclear accumulation of poly(A) RNA is an evolutionarily-conserved hallmark of quiescence across species and cell types.

**Fig. 5.**
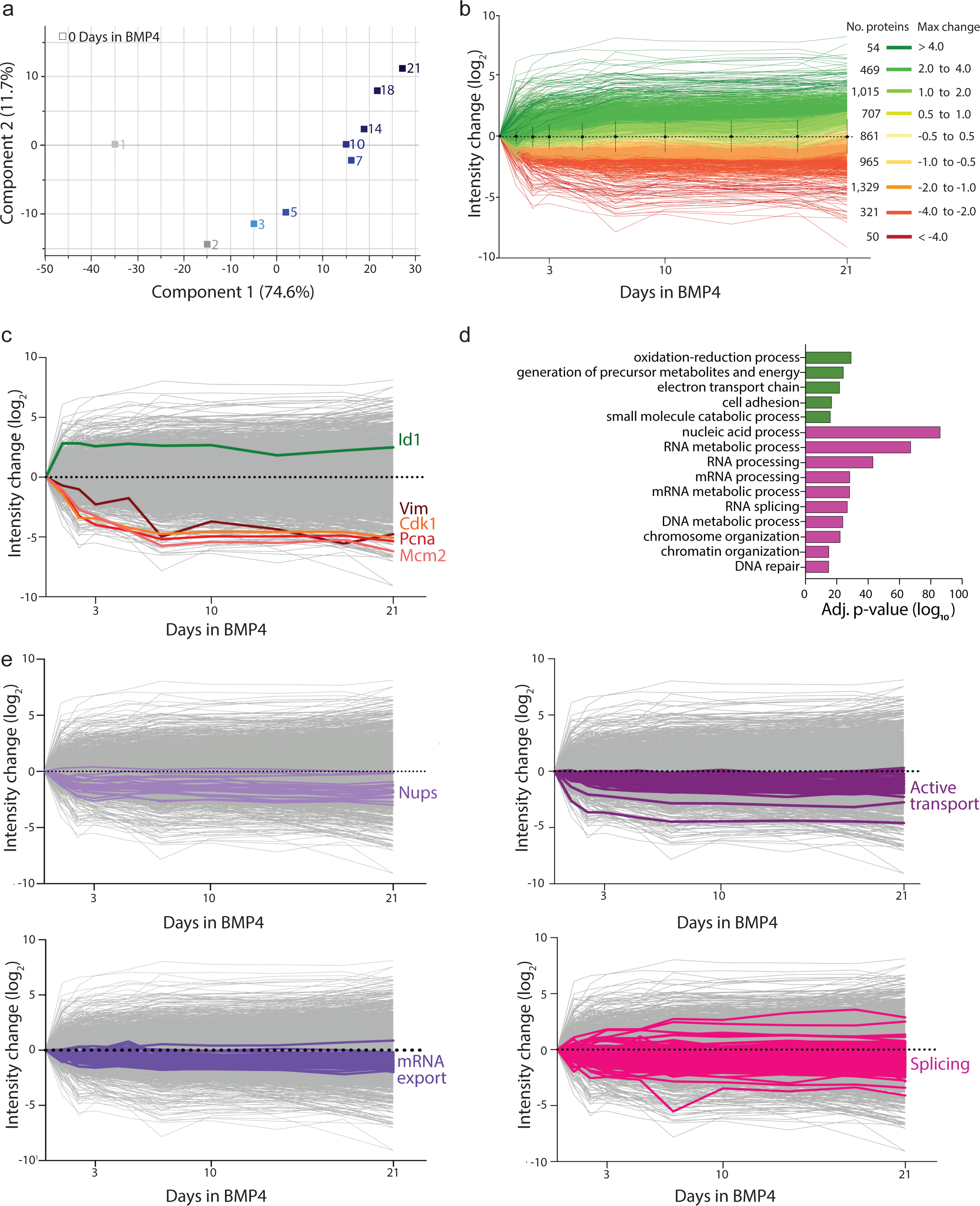
Proteome changes as NSCs transition between active and deeper quiescence states. **a**, PCA plot. b, TMT protein intensity changes (log_2_ difference, arbitrary units) from Day 0 (aNSCs); mean and S.D. indicated for each time-point.**c**, Control proteins plotted on background of all. **d**, Most upregulated (5, green) and down-regulated (10, magenta) biological process GO terms in 21d-qNSCs. **e**, Change of specified categories of proteins plotted on background of all.

## Downregulation of Nup98-96 in mouse NSCs induces quiescence features

To enquire into evolutionary conservation of the mechanism we found underpinning nuclear accumulation of poly(A) in quiescence in *Drosophila* NSCs, we knocked down Nup98-96 in mouse hippocampal NSC cultures. After confirming that Nup98 levels decreased in response to short-hairpin RNA (shRNA), we determined the proportion of cells positive for the proliferation antigen Ki67 and measured accumulation of nuclear poly(A). Nup98-96 knockdown samples had a reduced percentage of Ki67^+^ cells and accumulated nuclear poly(A) RNA in the absence of cell death as reported by cleaved Caspase 3 (Fig. 4f). This was observed with two independent shRNAs and the effect was reversible (Extended Data Fig. 6). We concluded that, like in *Drosophila* NSCs, attenuation of Nup98-96 in mouse NSCs induced quiescence.

**Fig. 6.**
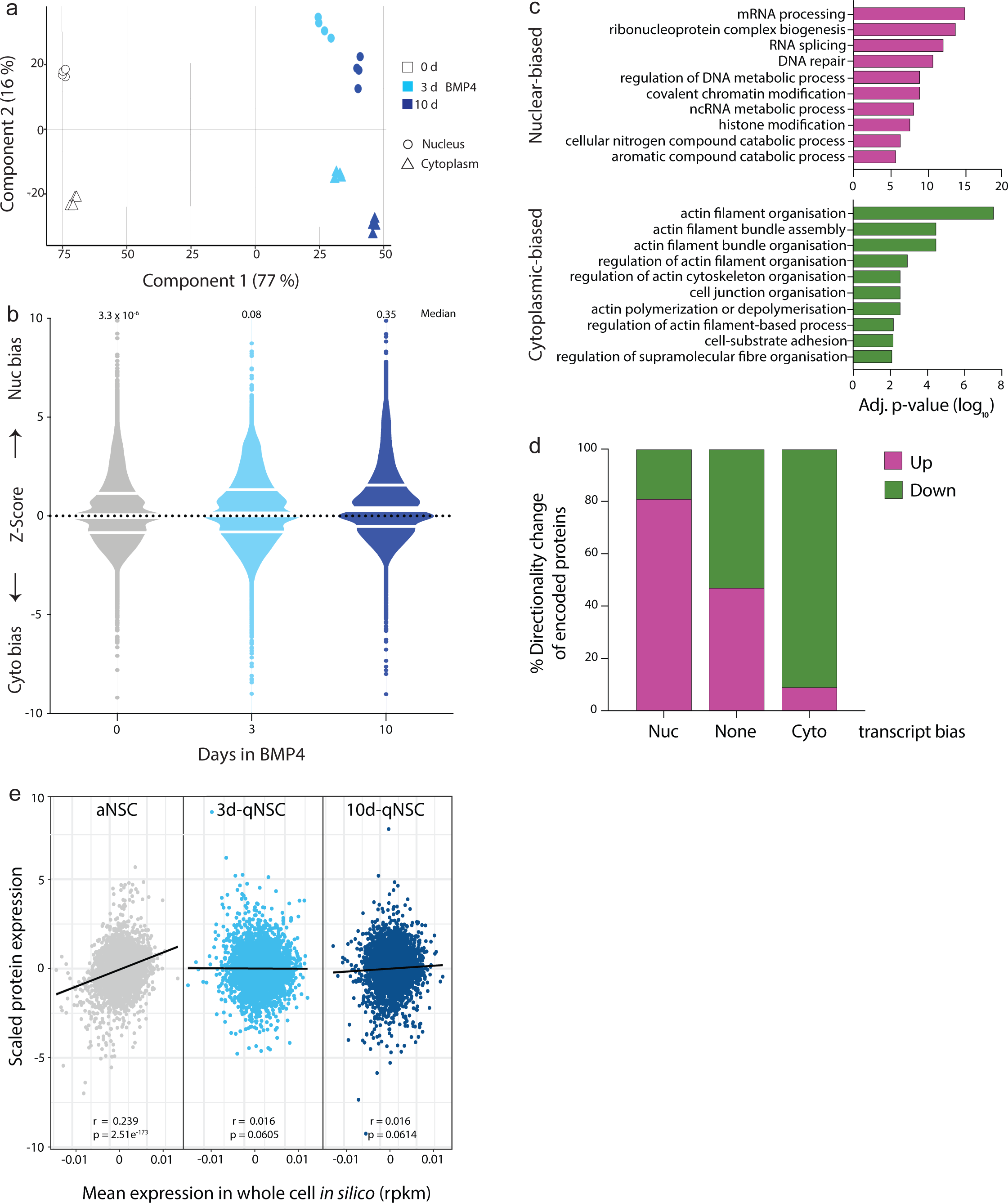
Nuclear retention of transcripts increases as NSCs transition between active and deeper quiescence states. **a**, PCA plot. **b**, Z-scores at 0, 3 and 10 d in BMP4 for the transcripts detected across all conditions (omitting eleven data points out of range); white lines: median and quartile boundaries. **c**, Most nuclear-biased (10, magenta) and cyto-plasmic-biased (10, green) biological process GO terms in qNSCs relative to aNSCs. **d**, Nuclear-or cytoplasmic-biased transcripts predict directionality of protein levels change in ∼80 % cases (shown correlation at 10 versus 0 d BMP4). **e**, The proteome becomes uncoupled from the transcriptome in qNSCs Mean expression in whole cell *in silico* (rpkm) (protein intensities scaled to normalise range of expression between samples).

## Most nucleocytoplasmic transport factors are downregulated in qNSCs

Whilst downregulation of nucleocytoplasmic transport components through experimental manipulation can induce NSC quiescence, we wondered whether this occurred physiologically. To determine whether nucleocytoplasmic transport factors are endogenously downregulated in qNSCs, we took advantage of the mouse NSC monoculture system to compare global proteome expression between aNSCs and qNSCs, not reported before. Protein extracts from adult mouse hippocampal NSCs were prepared on different days post-BMP4 addition (0 days being the active condition), having ascertained that longer exposure to BMP4 corresponded to deeper quiescence (Extended Data Fig. 7), and longitudinal proteome profiling was carried out by tandem mass tag (TMT) spectrometry^53^. Principal component analysis (PCA) showed that BMP4 exposure length accounted for the majority of protein changes already within 3 days (Fig. 5a), our standard quiescent condition. 5,771 proteins were reliably identified and quantified in all fractions, with over four-fifths changing considerably in either direction and nearly as many upregulated as downregulated over 21 days of BMP4 exposure (Fig. 5b and Supplementary Table 3). Validating the data, proteins whose levels are known to be altered in qNSCs behaved as expected^52, 54^ (Fig. 5c). Furthermore, the negative regulators of the TOR pathway Tuberous sclerosis proteins 1 and 2 (Tsc1 and Tsc2) were upregulated, and proteins involved in DNA replication and cell-cycle progression were well represented amongst those downregulated in qNSCs relative to aNSCs (Supplementary Table 3).

**Fig. 7.**
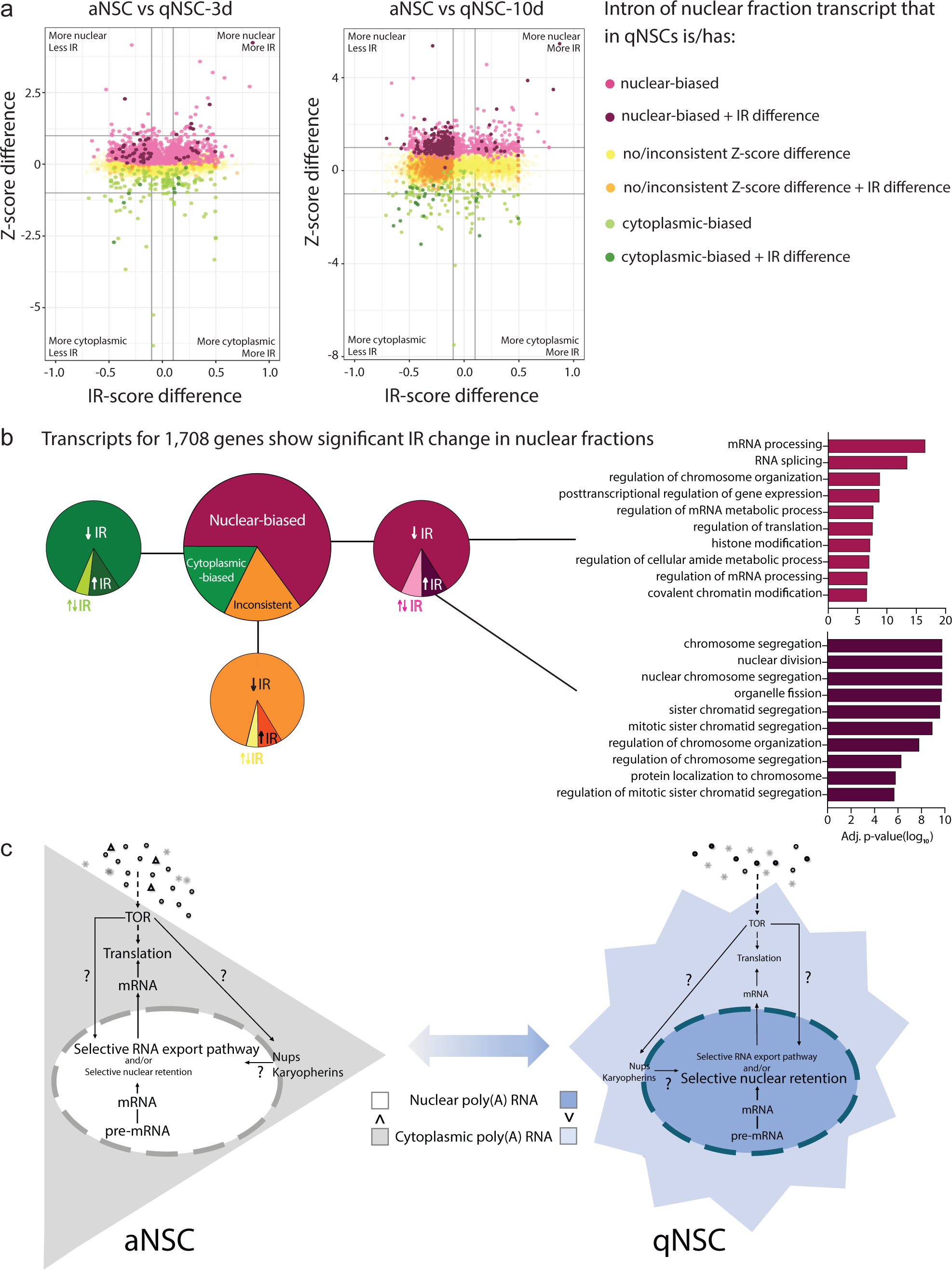
Nuclear-biased transcripts in qNSCs are generally more completely processed than in aNSCs. **a,b** depict data from nuclear fractions. **a**, Z-score and IR-score differences between aNSCs and qNSCs per gene (single intron with largest IR difference plotted). **b**, Large pie chart: nuclear fraction genes according to Z-score bias in qNSCs; small pie charts: (un)biased genes according to direction of differential IR. Right: most enriched biological process GO terms for genes in categories indicated. **c**, Model. A variety of qualitatively and/or quantitatively distinct signals converge on the TOR pathway to modulate cell activity or quiescence. This regulates translation via known targets but potentially also via the paths suggested with question marks towards nuclear retention of mRNA. Alternatively, the additional paths are controlled in a TOR-independent manner. Relative font size suggests degree of activity of various processes in aNSCs versus qNSCs.

Biological processes GO term analysis revealed the most downregulated categories of proteins in qNSCs as (m)RNA processing and metabolism, as well as DNA organisation and metabolism (Fig. 5d, Supplementary Table 4). Consistent with the hypothesis that poly(A) RNAs accumulated in the nucleus due to reduced availability of nucleocytoplasmic transport factors, we detected lowering levels of most of these as NSCs transitioned from an active state into deep quiescence; none were upregulated (Fig. 5e, Supplementary Table 3). Specifically, three-quarters of nucleoporins (23, containing members of all structural classes), were detected in our proteome-wide investigation, with 19 (representing all structural classes) consistently decreased in qNSCs, and 4 expressed at comparable levels across conditions (Fig. 5e, Supplementary Table 3). Employing independent methods (Western blots or immunocytochemistry), we confirmed significant downregulation of a few Nucleoporins after 3 days of BMP4 (Extended Data Fig. 8). This data suggests altered nucleoporin stoichiometry in qNSCs versus aNSCs although it remains unclear how it changes specifically at the NPC. Regarding active nucleocytoplasmic transport regulators, we detected 26, with levels of all but three decreasing as NSCs shifted into quiescence (Fig. 5e, Supplementary Table 3). In all, multiple nucleocytoplasmic protein transport factors are downregulated in qNSCs relative to aNSCs.

## Most known mRNA export pathway and splicing factors are also downregulated in qNSCs

mRNA export from the nucleus is coupled to pre-mRNA processing, including a checkpoint for completed splicing^31, 35, 37, 55^. Furthermore, and in line with proteomic downregulation of (m)RNA processing and metabolism components, we considered the possibility that the effect of altered nucleocytoplasmic transport on nuclear accumulation of poly(A) RNA in qNSCs might be due to limitation of mRNA export pathway components, upregulation of nuclear retention factors and/or incomplete splicing^37, 39^.

A few nucleoporins such as Nup96, Gle1, Nup155 and Tpr play key roles in mRNA export^37, 39^; the latter two were detected and both downregulated (Supplementary Table 3). We detected 38 mRNA export factors^37, 39^ of which 37 were downregulated in qNSCs (Fig. 5e, Supplementary Table 3). Concerning nuclear retention factors, only few are known^39^, and those picked up in our proteome analysis were downregulated in qNSCs, suggesting they are not responsible for the observed nuclear retention of poly(A) RNA (Supplementary Table 3). Regarding splicing factors, we detected 228 of which 204 were consistently downregulated as NSCs transitioned from active into deeper quiescence; intriguingly, 12 were consistently upregulated (Fig. 5e, Supplementary Table 3).

## Altered nucleocytoplasmic transcript bias uncouples proteome and transcriptome in qNSCs

To determine the identity and splicing status of nuclear-accumulated transcripts in qNSCs, we fractionated adult mouse hippocampal aNSC and qNSC cultures into nuclear and cytoplasmic compartments and performed RNA-seq on each (fracRNA-seq). We prepared samples from two quiescence depths, corresponding to 3 and 10 days BMP4 exposure (qNSC-3d and qNSC-10d). Transcripts corresponding to a total of 30,265 genes were detected, of which 19,979 (66 %) were protein-coding (Supplementary Table 5); about 95.5 % of reads mapped to protein-coding genes (Gene Expression Omnibus GSE162047). PCA showed progressive divergence of BMP4-treated NSCs from aNSCs as a function of time and further clustering of samples according to subcellular compartment (Fig. 6a).

For transcripts pertaining to each gene, and irrespectively of levels or splicing, we determined the proportion of exonic reads found in nuclear versus cytoplasm from paired extracts for each sample. For each gene we ascribed a bias score Z = log2 [number of nuclear reads / number of cytoplasmic reads], with Z > 0 indicating bias towards the nucleus and Z < 0 bias towards the cytoplasm (Fig. 6b, Supplementary Table 5). Validating the data, transcripts predicted to be biased towards nuclear or cytoplasmic compartments^56^ behaved as expected in all conditions and samples: *Nuclear Paraspeckle Assembly Transcript 1*, *Metastasis Associated Lung Adenocarcinoma Transcript 1* and *Plasmacytoma Variant Translocation 1* with a positive Z, indicating nuclear bias; *Ribosomal protein small subunit S14*, *Glyceraldehyde-3-phosphate dehydrogenase* and *Rn7s1* with a negative Z, indicating cytoplasmic bias.

In aNSCs, transcripts from 12,096 genes were significantly biased to either subcellular fraction, 5,637 (47 %) of which were nuclear-biased (3,026 with Z > 2 i.e., more than 4-fold bias). In qNSCs-3d, products of 12,812 genes were significantly biased, 6,601 (52 %) of which were nuclear-biased (3,470 with Z > 2). In qNSCs-10d, products of 12,227 genes were significantly biased, 7,653 (63 %) of which were nuclear-biased (4,106 with Z > 2). In a second step, we selected those genes whose products were detected in all conditions and performed pairwise comparisons between individual genes across conditions. Significant changes comprised genes whose products swapped bias in subcellular compartment as well as those whose Z scores changed significantly even if retaining the same compartment bias. Transcripts corresponding to 388 genes changed significantly between aNSCs and qNSCs-3d, of which 247 (64 %) became more nuclear-biased (20 more than 4-fold); 2,584 changed significantly between aNSCs and qNSCs-10d, of which 2,409 (93 %) became more nuclear-biased (311 more than 4-fold); and 590 changed significantly between qNSCs-3d and qNSCs-10d, of which 571 (97 %) became more nuclear-biased (128 by more than 4-fold) (Fig. 6b, Supplementary Table 5). Transcripts of 2,173 genes (86 % of which were protein-coding) changed subcellular bias score significantly and consistently (in the same direction between aNSCs and qNSCs-3d, then again between qNSCs-3d and qNSCs-10d) upon quiescence induction. Of these, transcripts of 1,616 genes (74 %; of which 92 % were protein-coding), became increasingly nuclear with quiescence. In summary, transcripts for most genes had no significant subcellular bias in any of the conditions nor, if they had, did the direction or magnitude of their bias change significantly between active and quiescent states. Notwithstanding, transcripts of more than 2,000 genes, mostly protein-coding, did change subcellular bias significantly, three-quarters of which becoming increasingly nuclear as NSCs shifted from active into deeper quiescence.

Concordant with proteomic downregulation, biological processes GO term analysis revealed the most nuclear-biased categories of transcripts in qNSCs as (m)RNA processing and RNP biogenesis, as well as DNA organisation and metabolism (Fig. 6c, Supplementary Table 6). Cytoplasmic-biased transcripts in qNSCs were enriched in actin cytoskeleton organisation GO terms, consistent with our observations of increased morphological complexity upon quiescence induction (Extended Data Fig. 9 and seen in flies by fibre extension) and enriched “cell adhesion” GO term in proteomics. Moreover, we observed ∼80 % concordance between increased nuclear-and cytoplasmic-bias of transcripts and respective protein down-or upregulation in qNSCs (Fig. 6d). This reveals a major impact of subcellular partitioning of transcripts on directionality of protein levels in qNSCs, explaining our observation that transcriptome and proteome become uncoupled in qNSCs (Fig. 6e). We conclude that altered nucleocytoplasmic distribution of transcripts in qNSCs contributes greatly to regulation of protein expression.

## Nuclear-biased transcripts are generally more completely processed in qNSCs than in aNSCs

To assess whether transcript nuclear-bias during quiescence could be globally accounted for by decreased splicing, we analysed intron retention (IR). For each intron of transcripts found in each subcellular fraction we ascribed an IR score = [number of intronic reads / (number of intronic reads + number of spliced reads)], with 0 < IR < 1. The majority of significant IR changes were observed in nuclear fractions as might be expected, with those in cytoplasmic fractions an order of magnitude lower (Extended Data Fig. 10a, Supplementary Table 7). In nuclear fractions, 3,362 introns corresponding to 1,708 genes showed significant IR changes between at least two conditions (Fig. 7a, Supplementary Table 7). Of these, 1,110 (65 %) genes corresponded to transcripts that became increasingly nuclear in qNSCs, 302 (18 %) to ones that became increasingly cytoplasmic, and 296 (17 %) to ones that showed no consistent subcellular change (Fig. 7b pie charts, Supplementary Table 7). For the most part, the various introns of a transcript showed the same directionality of IR as NSCs shifted from active into deeper quiescence (Extended Data Fig. 10c, Supplementary Table 7) and the vast majority of significant IR changes in the nucleus consisted of IR *decrease*.

In agreement with the categories of transcripts that became increasingly nuclear as quiescence deepened, biological process GO terms for the 934 genes encoding nuclear-biased transcripts with decreased IR in nuclear fractions pertained to (m)RNA processing and chromatin regulation; and those for the 98 nuclear-biased transcripts with increased IR pertained mostly to mitosis, followed by (m)RNA processing (Fig. 7b right, Supplementary Table 8). In summary, despite a small and potentially meaningful category of genes where increased IR positively correlates nuclear-biased transcripts, most transcripts that become more nuclear during NSC quiescence are not retained in the nucleus due to increased IR and are in fact more completely spliced.

A recent study reported widespread IR in various quiescent cell types^57^ but did not report on qNSCs nor did we find this in our study. It is possible that different cell types adopt distinct molecular strategies towards the same goal of selective nuclear-bias of transcripts. Indeed, we report increased nuclear-to-cytoplasmic ratio of poly(A) RNA in a few quiescent cell types relative to active counterparts, across three different organisms. Nuclear retention of mRNAs has been observed in reaction to stress or changing cellular conditions, such as differentiation, viral infection or oncogenic transformation. Nuclear retention protects mRNAs from viral nucleases and cytoplasmic decay pathways whilst release allows rapid cellular responses^39^.

Studies in human fibroblasts and zebrafish NSCs have reported Exportin-1-dependent changes in microRNA biogenesis and localisation in quiescence, including accumulation of microRNA-9 and Argonaute proteins (components of RNA silencing complexes) in qNSC nuclei^55, 59^. In *Drosophila*, a transient low-level nuclear pulse of the homeobox transcription factor Prospero (Pros) induces *Drosophila* NSC quiescence^60^ (its localisation matching that of its adaptor Mira, shown in Extended Data Fig. 1). RanGEF/Rcc1/Bj1 has been implicated in excluding Pros from aNSC nuclei to allow their self-renewal^61^ and its downregulation might therefore enable quiescence-inducing nuclear Pros. The work here presented shows these to be glimpses into larger-scale nucleocytoplasmic transport alterations that control quiescence. We show that downregulation of various nucleoporins or active nucleocytoplasmic transport factors induce NSC quiescence; and that many of these proteins are downregulated in response to a quiescence signal. Furthermore, as expected from altered nucleocytoplasmic transport, we identify cargo that are differentially partitioned between nucleus and cytoplasm in active versus quiescent NSCs. Specifically, we demonstrate nuclear-bias of hundreds of transcripts presumed to orchestrate a finely-tuned temporal sequence of events in the regulation of quiescence.

Nuclear-biased mRNAs with decreased IR mostly encode (m)RNA processing regulator whereas those with increased IR mostly encode cell-cycle and mitotic regulators. These mechanisms add nuance to downregulation of protein synthesis during quiescence, via sequential deployment of factors during quiescence entry and, presumably the reciprocal during quiescence exit. Gene products relying differentially on transcription, splicing and/or cytoplasmic translocation will need more or less time for deployment and the first will likely promote readiness of others to follow. Both towards quiescence entry and exit, the post-transcriptional mechanisms described likely interplay with the transcriptional in a series of positive-feedback loops whereby initially small changes in protein or cytoplasmic transcripts are amplified, underlying the continuum of states between deep quiescence and active cells. In fact, the same components may in some cases regulate quiescence at multiple levels and even different mechanisms. Analyses of *Drosophila* mutants for the nucleoporin Mtor/Tpr (whose RNAis did not induce quiescence in this study) implicated it in NSC quiescence regulation via transcriptional effects of the so-called spindle matrix complex^62^.

The course of transcription, transcript processing, export and degradation results in a steady-state that determines nucleocytoplasmic partitioning of transcripts. Our analyses reveal that most NSC mRNAs are equilibrated between the two subcellular compartments but that transcripts for a few hundred genes become more nuclear (and more completely spliced) in qNSCs. The fact that subcellular compartmentalisation of most transcripts was not significantly altered between aNSCs and qNSCs argues against general disruption of splicing or mRNA export. It is possible that nuclear-biased transcripts benefit especially from the few splicing factors that exhibit higher protein levels during quiescence (Fig. 5e). Another possibility is that overall decrease in splicing is commensurate with decrease in transcriptional rate during quiescence, and that decrease in IR results from longer residence of nuclear-biased transcripts in the nucleus allowing sustained interactions with splicing factors towards completion. Since the bulk mRNA export pathway is non-discriminatory and that its canonical components were only modestly downregulated in quiescence, it is plausible that qNSCs selectively regulate (a) discriminatory mRNA nuclear retention and/or export pathway(s) of a few hundred transcripts. Selective mRNA export pathways coordinate export of functionally related mRNAs^37, 39^, which we see among nuclear-biased transcripts. However, neither are all components of selective pathways currently known nor is the complement of mRNAs exported by each known pathway identified^37^. Future work will uncover (the) mechanism(s) for nuclear-bias of transcripts in qNSCs (Fig. 7c).

Taken together, our results establish several novel features of NSC quiescence regulation, at least some of which may prove generalisable to other quiescent cell types. First, aNSCs and qNSCs express different concentrations and stoichiometry of nucleoporins, karyopherins and mRNA export factors; second, perturbation of specific components of the nucleocytoplasmic transport machinery is sufficient to induce quiescence; third, induction of qNSCs by physiological cues or via nucleoporin downregulation causes nuclear accumulation of a susbtantial fraction of polyadenylated transcripts and the magnitude of this effect increases with quiescence depth; fourth, nuclear or cytoplasmic biases of transcripts largely predicts down-and upregulation of encoded proteins, respectively, evidencing impact of mRNA nuclear accumulation on the proteome of qNSCs and explaining our finding of uncoupled proteome and transcriptome in qNSCs; fifth, most nuclear-biased mRNAs are completely spliced and encode (m)RNA processing regulators whereas the minority with increased intron-retention mostly encode cell-cycle and mitotic regulators; this suggests sequential deployment of types of factors during quiescence entry and, presumably the reciprocal during quiescence exit.

The study of quiescence at both fundamental and clinical levels has been hampered by lack of positive markers. By showing that quiescence uncouples the proteome from the transcriptome, our study sheds light on why markers of quiescence may not have emerged from transcriptomic studies, and opens new avenues to identify them. Our findings also unveil likely mechanisms underlying graded states of stem cell quiescence and the rapid return of quiescent cells into the cycling pool upon receiving appropriate cues.

## Methods

### Drosophila melanogaster genetics

Flies were mutagenised and screened as published^63, 64^. MARCM clones^65^ were induced as described^64^ using *y,w,hs-FLP^1.22^;tub-GAL4,UAS-NLS::GFP::6xmyc;FRT82B,tubP-GAL80^LL3^/(TM6B)* (gift from G. Struhl).

RNAi was performed in conjunction with *UAS-Dcr2*^66^ and stocks were obtained from the Transgenic RNAi Project at Harvard Medical School, Vienna *Drosophila* Resource Centre (VDRC), the Japanese National Institute of Genetics (NIG) and Bloomington *Drosophila* Resource Center (BDRC); deficiency^67, 68^ and balancer stocks, *UAS-mCD8::GFP*, *UAS-Dcr2 and w; tub-GAL80^ts^* were also obtained from the BDRC. The following additional strains were used: *grh-GAL4*^18^ and *UAS-mira::3xGFP*^64^ to visualise a subset of NSCs; *NP3537-GAL4*^78^ (NIG) for pan-NSC RNAi.

### Rearing and staging of *Drosophila*

For larval genotyping, lethal chromosomes were re-established over balancer chromosomes marked by Dfd-YFP. For larval staging experiments, crosses were performed in cages with grape juice plates (25 % (v/v) grape juice, 1.25 % (w/v) sucrose, 2.5 % (w/v) agar) supplemented with live yeast paste. Larvae hatched within 2 h at 25°C were transferred to our standard cornmeal food (8 % (w/v) glucose, 2 % (w/v) cornmeal, 5 % (w/v) baker’s yeast, 0.8 % (w/v) agar in water) and placed at the desired temperature. Early larvae were newly-hatched and late larvae were at wandering stage. Data from males or females of the same genotype were pooled without distinction.

### Tissue/cell collection, and cell culture

*Drosophila* or mouse tissue/cells were prepared and cultured as published^52, 63, 70^. Differences were that for mouse cells the medium for aNSCs was additionally supplemented with 20 ng/ml recombinant murine Epidermal Growth Factor (PeproTech 315-09) and qNSC induction was performed with 50 ng/ml BMP4 (R&D Systems 5020-BP-010). qNSCs cultures were never passaged. *Nestin-GFP* mice^71^ (n = 3) were used for *in vivo* NSC analysis. Umbilical cord blood derived HSCs were extracted and processed as previous published^72^. Immunophentoypic HSCs were defined as Lin^-^/CD34^+^/CD38^-^/CD45RA^-^/CD90^+^/CD49f^+^, where Lin is Human Lineage Cocktail 1 (CD3, CD14, CD16, CD19, CD20, CD56); sorted by an Aria cytometer (BD Biosciences). For all flow cytometry, cells were initially identified based on forward and side scatters. Dead cells were excluded based on staining with 4’,6-diamidino-2-phenylindole (DAPI).

### EdU labelling, oligo(dT) fluorescent *in situ* hybridisation (FISH) and Immunofluorescence

Tissue/cells were incubated in 10 mM EdU (ThermoFisher C10340) in medium (*Drosophila* CNSs for 2 h in Schneider’s Insect Medium; mouse NSCs for 6 h in aNSC or qNSC media described above). Samples were processed for EdU detection according to the manufacturer’s instructions (ThermoFisher Click-iT EdU Imaging Kit). For combined EdU and other stains these were performed before EdU colour reaction. For FISH of *Drosophila* tissue or mouse cultured cells, samples fixed as usual were permeabilised in cold Methanol for 10 min, rehydrated in 70 % Ethanol for at least 10 min followed by 1 M Tris-hydroxymethyl-aminomethane (Tris) buffer (pH 8.0) for 5 min. For FISH of mouse tissue, 40 µm coronal sections first underwent heat-mediated antigen retrieval at 95 °C for 10 min in 10 mM Saline-Sodium Citrate buffer (SSC) (pH 6.0). For all samples, hybridisation was performed with 1 ng/μl Cy3-Oligo-dT(50) (Genelink, 26-4322-02) in 2x SSC containing 1 mg/ml yeast tRNA, 0.005 % Bovine Serum Albumin, 10 % Dextran Sulfate and 25 % deionised Formamide; for at least 2 h at 37 °C in a humidified chamber. Samples were washed once in 4x SSC and twice in 2x SSC. For combined FISH and immunofluorescence, FISH was performed before primary antibody incubation in 2x SSC, 0.1 % TritonX-100 (with 5 % donkey serum for mouse tissue, all other samples, no serum) and subsequent steps in 2x SSC. When performing immunofluorescence alone or with EdU, this was performed as published^19, 52, 63, 70^ with additional primary antibodies: rabbit anti-Dcp-1 (Cell Signalling 95785), rabbit anti-cleaved Caspase-3 (Cell Signalling Technology 9664), rabbit anti-Nup98 (Cell Signalling Technology 2598), rabbit anti-Nup214 (Abbex abx129466), rabbit anti-Nup54 (Novus NBP1-85899), mouse anti-Nup133 (Novus H00055746-M02), rabbit anti-Nup43 (Novus NBP1-88792), rabbit anti-Ndc1 (Novus NBP1-91603). Antibodies used for fluorescence-activated cell sorting were: anti-Human Lineage Cocktail (BD Biosciences 340546), anti-CD34 (BD Biosciences 8G12), anti-CD38 (eBioscience HB7), anti-CD45 (Biolegend HI30), hCD45RA (eBioscience HI100), anti-CD90 (BD Biosciences 5E10), anti-CD49f (BD Biosciences GoH3).

### Imaging and image analyses

Fluorescence samples were scanned with Zeiss 510 or 800, or Leica SP5 or SP8 scanning confocal microscopes. Optical section steps ranged from 0.1 to 2 μm with picture size of 1,024 × 1,024 pixels. Images were processed and arranged using Fiji/ImageJ, Adobe Illustrator, Adobe Photoshop CS5, and/or PowerPoint software. *Drosophila* cell counts were carried out with ImageJ Cell Counter plugin; when qNSC fibres were observed 10 brain lobes were quantified per genotype, when not observed it was 4 lobes (a single lobe per animal). Cell culture images were acquired from 3 random fields from each of 3 coverslips; their counts, image masking, projecting and reformatting were performed with CellProfiler scripts. Poly(dT) intensity per area was determined for nucleus or cytoplasm from SUM projections when the whole cell could be imaged, or from single optical sections of mouse hippocampus (where dense cell packing and sectioning precluded imaging entire cells). No data points were excluded. Graphpad Prism software was used for statistics and graphs.

### Lentivirus preparation and titration

Lentiviruses were produced in 293FT using standard procedures and titratred as published^73^. To deplete Nup98, NSCs were grown for 24 h, infected with 2.5 M.O.I. Nup98 shRNA lentivirus and fixed 0 or 2 d post-infection. To reversibly induce shRNA-mediated knockdown, the medium was supplemented with 1 ! g/ml Doxycycline hyclate (Sigma D9891) daily; reversed by washing three times with basal medium.

### Protein extraction, Western blots and proteomics

Cells were washed with ice-cold PBS and scraped. Protein lysates were prepared as published^52^ and concentration determined using Pierce BCA Protein Assay kit (ThermoFischer Scientific 23225). Western blots were performed and quantified as published^52^. Antibodies used were all from Novus: rabbit anti-Nup50 (NBP2-19610), rabbit anti-Nup210 (NB100-93336), rabbit anti-Nup93 (NBP1-81546), rabbit anti-Nup62 (NBP1-31381), rabbit anti-Nup35 (NB100-93322), rabbit anti-Nup188 (NBP1-28717), mouse anti-Nup358 (NB100-74480), rabbit anti-Pom121 (NBP2-19890). Proteomic analyses were performed on 50 µg total protein per timepoint. Samples were Acetone-precipitated overnight followed by Trypsin digestion using the PreOmics iST-NHS sample preparation kit^74^, labelled using 0.2 mg TMT 10-plex Isobaric Label Reagents (Thermo Scientific) and checked to ensure >99 % labelling efficiency. Equal volumes of all ten labelled samples were mixed to produce a single mixed sample which was subject to high pH (HpH) reversed-phase peptide fractionation (Pierce). Nine HpH fractions were each analysed on a 145 min U3000 HPLC method. Samples were loaded in 2 % Acetonitrile, 0.05 % Trifluoroacetic acid onto a C18 trap column, then transferred onto an EasySpray 50 cm × 75 µm column. Peptides were separated by elution using the following conditions: 15 min 3-9 % mobile phase A (0.1 % Formic acid, 5 % Dimethyl Sulfoxide (DMSO)), 90 min 9-30 %, 15 min 30-50 %, 5 min 99 % and ending with 15 min at 3 %. Mobile phase B was 80 % Acetonitrile, 5 % DMSO, 0.1 % Formic acid. An SPS-MS3 method on an Orbitrap Fusion Tribrid mass spectrometer (Thermo Fisher Scientific) acquired data with settings: MS1 orbitrap, resolution 120 K, scan range 375-1500 m/z, maximum injection time 50 ms, AGC target 4E5, normalized AGC target 100 %, microscans 1, RF lens 30 %, profile data, MIPS mode peptide, charge states 2-6 included, dynamic exclusion 60 s +/-10 ppm. MS2 ion trap, quadrupole isolation mode, 1.2 isolation window, CID activation, 35 % collision energy, activation time 10 ms, activation Q 0.25, turbo scan rate, maximum injection time 50 ms, AGC target 1E4, normalised AGC target 100 %, microscans 1, centroid data, filter precursor selection range MSn 400-1200 m/z. MS3 orbitrap scan event 1 for charge state 2, quadrupole isolation mode, 1.3 isolation window, Multi-notch Isolation True, MS2 Isolation Window (m/z) 2, number of Notches 5, activation type HCD, collision energy 65 %, orbitrap resolution 50K, scan range 100-500 m/z, maximum injection time 105 ms, AGC target 1E5, normalized AGC target 200 %, microscans 1, centroid data. MS3 orbitrap scan event 2 for charge state 3 as above but with number of Notches 10. MS3 orbitrap scan event 3 for charge states 4-6 as above but with number of Notches 10. Raw data were analysed in MaxQuant^75^ (v1.6.12.0) against a SwissProt *Mus musculus* protein database containing 17,482 protein entries (downloaded May 2020). TMT10plex quantification was selected (modification at Lysine and peptide N-terminal amino groups) along with variable modification of Methionine oxidation and N-terminal acetylation. A fixed Cysteine modification of +113.084 Da (specific to the iST-NHS kit) was added. Further data analyses were performed in Perseus^76^ (v1.4.0.2). Common contaminants and proteins identified from decoy sequences were removed. Protein intensities were log2 transformed, median normalised within each sample and then normalised to Day 0. GO terms were simplified using REVIGO^77^ with allowed similarity of 0.7.

### Cell proliferation analysis by flow cytometry

Fixed samples were analysed with a BD LSRFortessa^TM^ Flow Cytometer using 488 nm excitation (505 LP, 530/30) for zsGreen, 355 nm (-, 450/50) for DAPI, and 639 nm (-, 670/14) for AlexaFlour 647-labelled EdU. Results were analysed with FlowJo software.

### Cell fractionation, nuclei acid extraction, sequencing and analyses

*Drosophila* genomic DNA was extracted according to standard methods and the identity of the genetic lesion determined by sequencing exons of heterozygous *FRT82B/FRT82B, 2V327* animals. This sequencing was outsourced to Eurofins Genomics using primers designed with A Plasmid Editor software. Mouse cell nucleocytoplasmic fractionation and RNA isolation was performed with the PARIS kit (ThermoFisher AM1921) according to the manufacturer’s instructions plus an additional ethanol precipitation step and rehydrated in nuclease-free water. Fractionation quality was verified by quantitative reverse-transcriptase polymerase chain reaction as published^78^. Subsequent steps for next generation sequencing were performed as published^52^ and sequencing was performed with 75 bp single-end reads with a depth of 50 million reads per sample. Raw reads were quality and adapter trimmed using cutadapt-1.9.1 software^79^ then aligned and quantified using RSEM-1.3.0/STAR-2.5.2^80, 81^ against the mouse genome GRCm38 and annotation release 89, both from Ensembl. Differential gene expression analysis was performed in R-3.6.1 (R Core Team, 2019) using the DESeq2 package^82^ (version 1.24.0). Normalisation and variance-stabilising transformation was applied on raw counts before performing PCA and Euclidean distance-based clustering. Significantly differential genes were always selected using a 0.05 false-discovery rate threshold. Size factors in DESeq2 were calculated based on the summed cytoplasmic and nuclear counts for each paired sample set to reconcile technical differences between samples whilst count differences between subcellular compartments within the same paired sample set were maintained for all genes. We performed pairwise comparisons between subcellular compartments on each day (within day comparison) and between the subcellular distribution across days (between days) using the following formula: ∼ days_in_BMP + subcellular_compartment + days_in_BMP:subcellular_compartment + days_in_BMP:pair_within_day. Additionally, we performed a likelihood ratio test in DESeq2 to identify the genes that changed subcellular distribution across time (reduced design formula in DESeq2: ∼ days_in_BMP + subcellular_compartment + days_in_BMP:pair_within_day). Z-scores of protein-coding genes (from fracRNA-seq data) was correlated to respective protein expression (proteomic data) in R-3.6.1 (R Core Team, 2019) using dplyr package (version 1.0.6). Scatterplots were prepared using ggplot2 (version 3.2.1) to visualise IR as one intron per gene, selecting the one with the biggest absolute change in IR value. Violin plots were prepared using ggplot2 (version 3.2.1) using the log2 fold changes for all genes after pairwise comparisons between days. Gene set enrichment analyses were performed using ClusterProfiler (version 3.12.0^83^) using the enrichGO() function for enrichment of biological process gene ontology terms.

### Data Availability

FracRNA-seq data is deposited in the Gene Expression Omnibus data repository and available under GSE162047.

### Code Availability

Code is available upon request.

## Acknowledgements

We are grateful to W. G. Somers and W. Chia for their contributions at the screening stage of this project; to F. Oozeer; plus R. Goldstone and J. Nicod (The Francis Crick Institute Advanced Sequencing Facility) for technical support; to A. Brand, S. Bray, F. Matsuzaki, C. Petritsch and G. Struhl for flies or antibodies; to the TRiP (NIH/NIGMS R01-GM084947), VDRC, NIG and BDRC stock centres for fly stocks; to Q. Ch’ng, R. Faraway, C. Houart, L. Lin, G. Neves, P. Rigo and S. Ultanir for discussions or comments on the manuscript. Work in the F.G. laboratory is supported by The Francis Crick Institute, which receives its funding from Cancer Research UK (FC0010089), the UK Medical Research Council (FC0010089) and the Wellcome Trust (FC0010089). This work was funded by a Cancer Research UK Career Development Fellowship to R.S.N. (A14958).

## Author contributions

A.C., M.M., A.R., A.M, L.H., A.C., H.F., R.C., P.F. and R.S.N. performed and analysed experiments; W.G. and D.B. provided human HSCs; P.F. and A.P.S. analysed raw proteomic data; S.S. and G.K. analysed raw fracRNA-seq data; A.R. and L.H. performed further analyses on proteomic and fracRNA-seq; R.S.N. designed and supervised the project, some aspects of which in collaboration with F.G.; R.S.N. wrote the manuscript, which all authors provided input to or approved.

## Additional information

Correspondence and requests for materials should be addressed to.

## Extended Data

Extended data consists of 10 figures and 8 supplementary tables.

**Extended Data Fig. 1.**
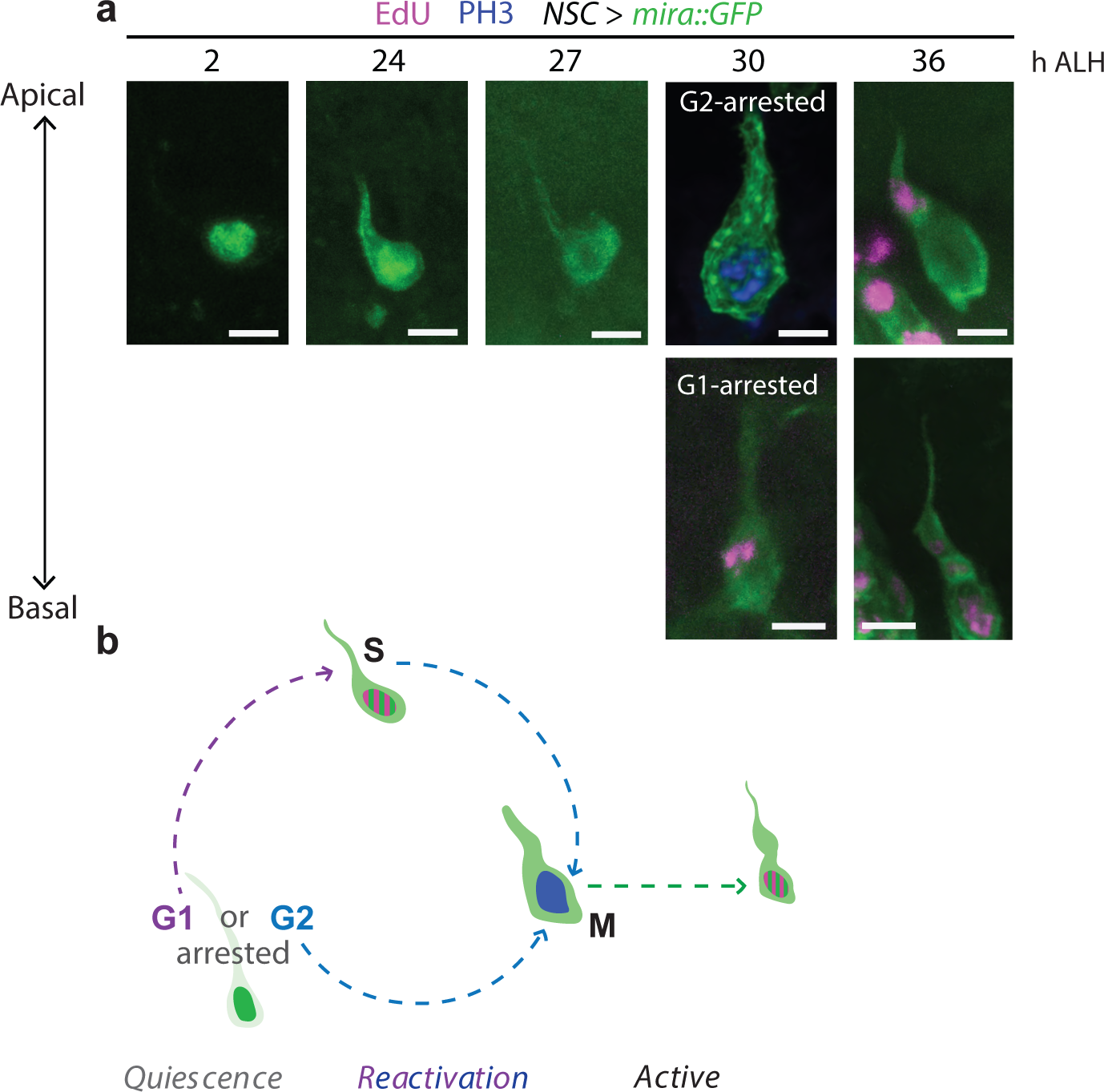
Sequence of events during *Drosophila* qNSC reactivation. **a**, Top row: timecourse of Mira::GFP localisation in a NSC reproducibly identified across specimen and used as model (ALH, after larval hatching; note that precise timings vary between NSCs). Deeply quiescent NSCs present nuclear Mira, a cell--body of ≤ 6µm and a long thin basal fibre (≤ 0.2 µm at the neck, i.e., junction with cell body). As NSCs emerge from deep quiescence, Mira localises not only to the nucleus but also to the cell cortex, decorating the fibre; the fibre thickens and eventually Mira is excluded from the nucleus. NSCs arrested in G2 such as this model, start expressing the mitotic marker phos-pho-histone H3 (PH3) without incorporating the S-phase marker 5-ethynyl-2’-deoxy-uridine (EdU). Bottom row (different timeline): G1-arrested NSCs incorporate EdU prior to enterting mitosis and becoming PH3--positive. Cell-cycle re-engagement markers are seen and mitosis completed whilst NSCs still harbour a fibre. Fibres are lost via inheritance by the firstborn post-reactivation basal daughter, a transit-amplifying progenitor named ganglion mother cell (GMC) in *Drosophila.* Images are maximum intensity projections of a few optical sections; scale bars: 5 µm. **b**, Schematic representation of these events colour-coded as per a.

**Extended Data Fig. 2.**
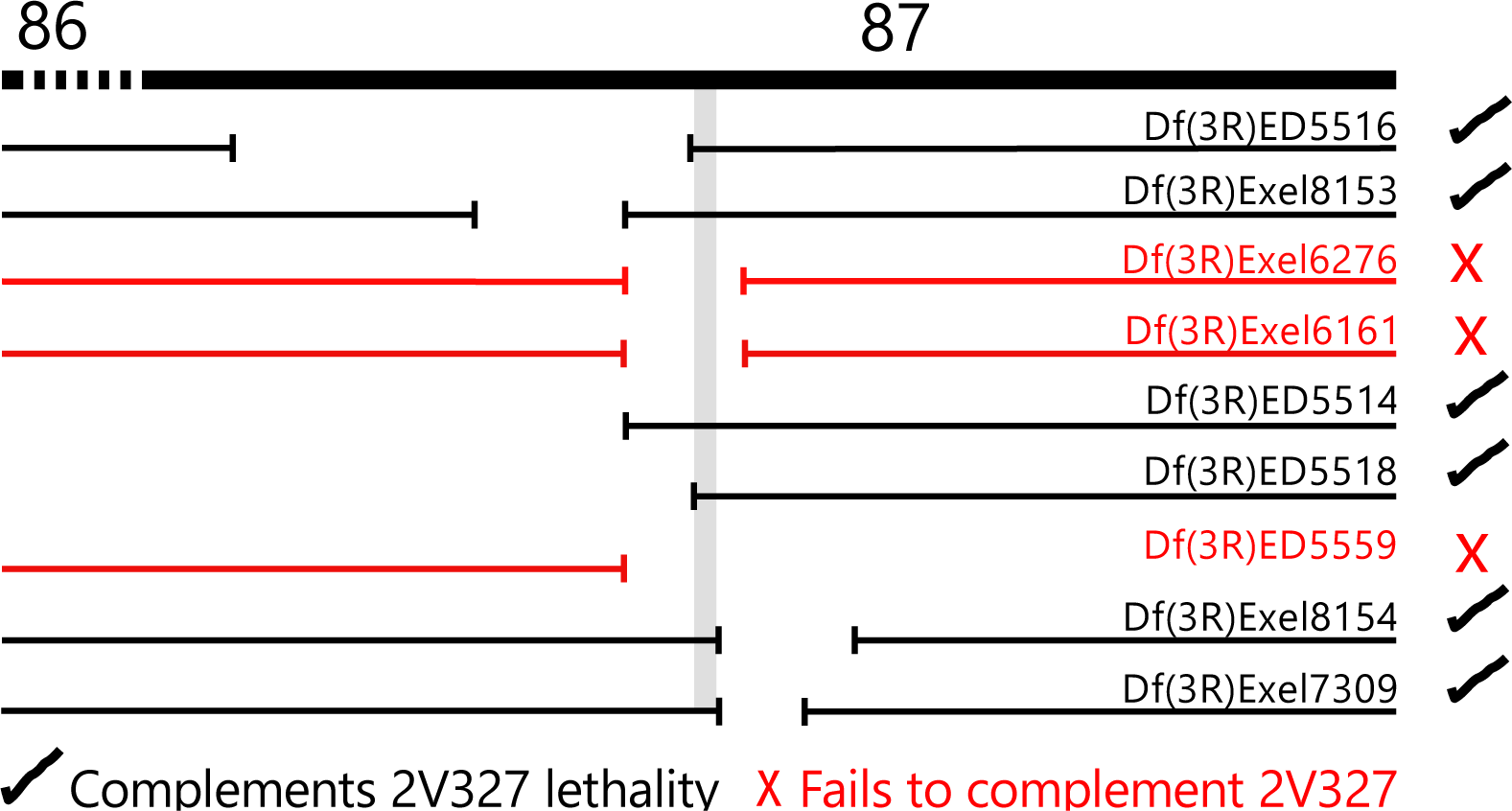
Deficiency mapping defined a small genomic interval responsible for the 2V327 phenotype. Phenotype in FRT82B MARCM clones revealed the 2V327 genomic lesion to be located on chromosome arm 3R. Assuming lethality of the mutation, complementation tests with DrosDel and Exelixis deficiencies (Df) uncovering 3R exposed a small candidate region between cytological locations 86-87 (grey). Regions uncovered by each Df are indicated by the gap in the line under each Df name.

**Extended Data Fig. 3.**
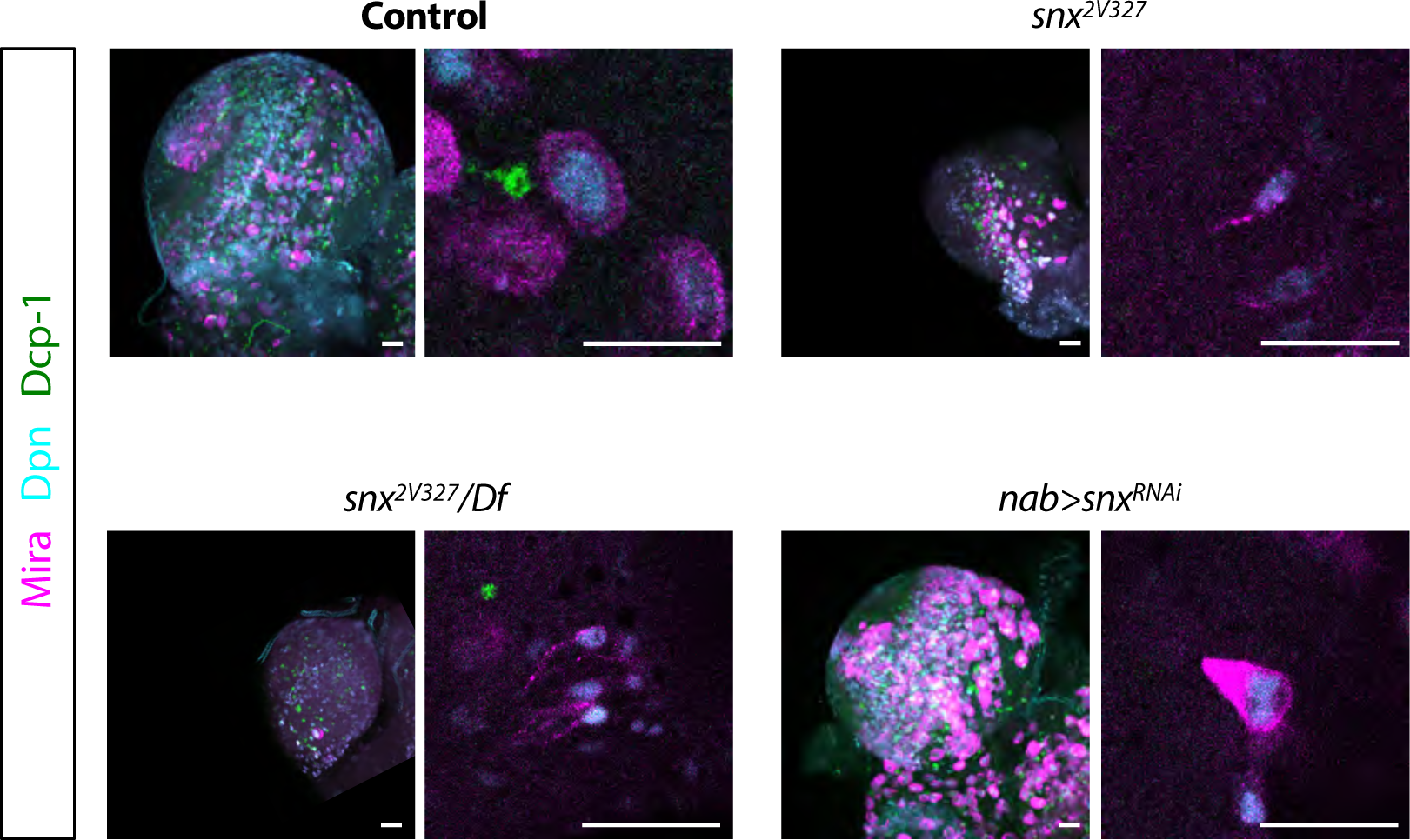
Mutant or RNAi *snx* NSCs do not express detectable levels of cleaved *Drosophila* Death Caspase-1. Shown is a low magnification of a whole late larval *Drosophila* brain lobe (left) and a high magnification of one/few NSCs (right) of the genotypes indicated. Scale bars: 15 µm.

**Extended Data Fig. 4.**
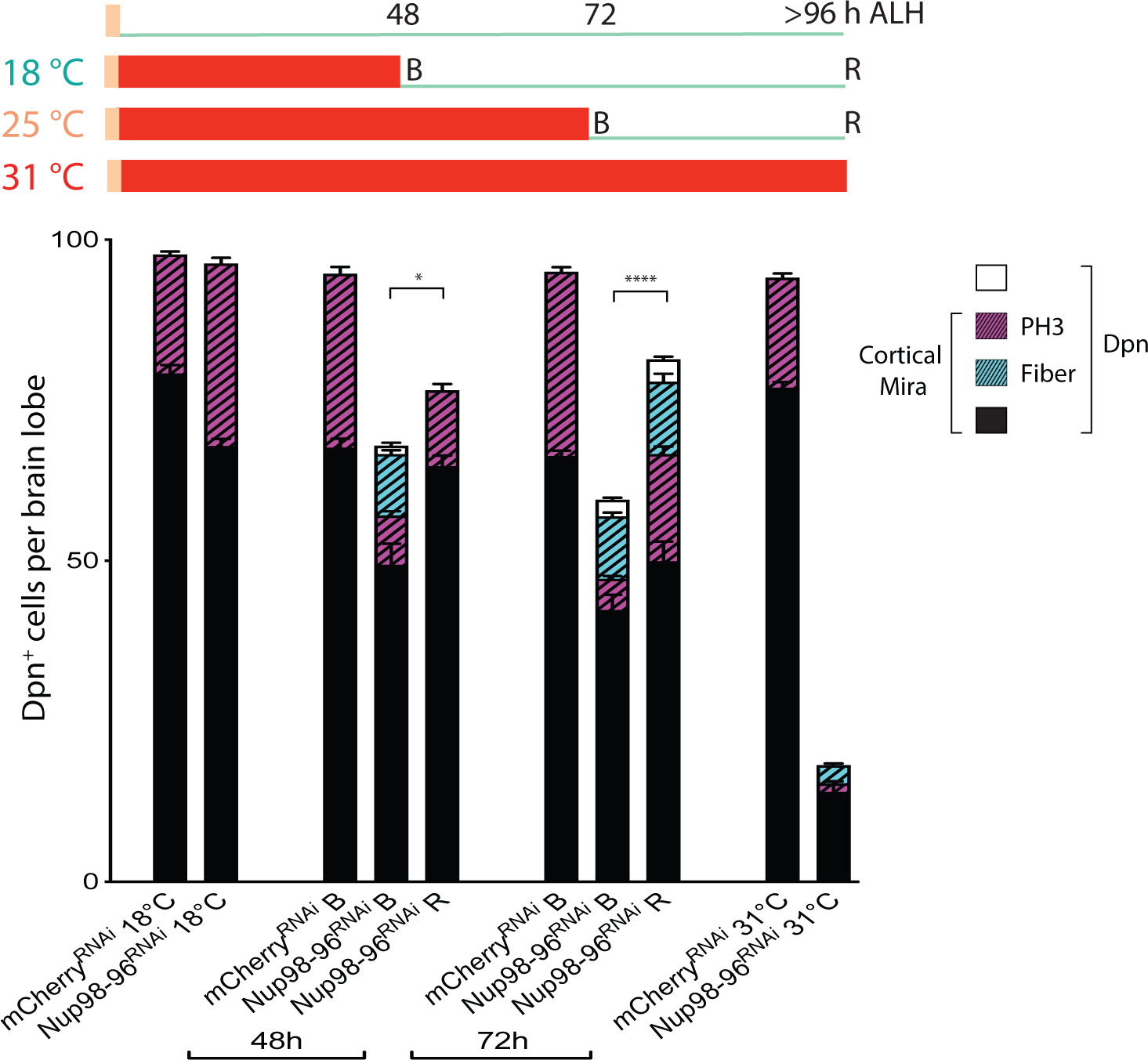
Downregulation of *Drosophila* Nup98-96 induces anachronical qNSCs. Experimental design (schematised) and quantifications as per Fig. 1d,e. Histograms represent mean and error bars s.e.m. Student’s t-test was performed to compare number of Dpn^+^ cells in the conditions indicated; *p<0.05. ****p<0.001.

**Extended Data Fig. 5.**
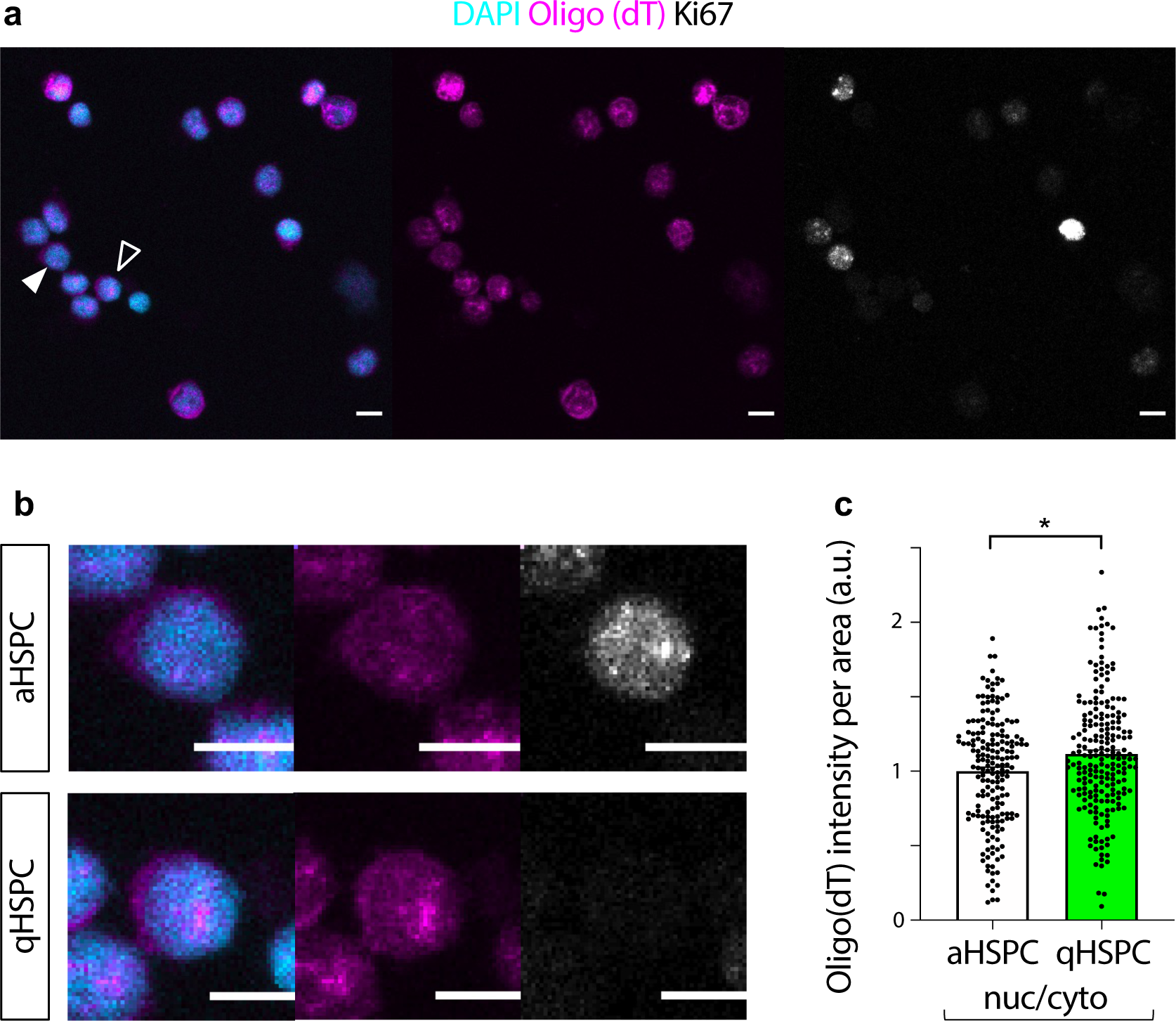
Quiescent haematopoietic stem cells accumulate nuclear poly(A) RNA relative to active counterparts. **a,** Adult bone marrow derived CD34+CD38-haematopoietic stem and progenitor cells (HSPCs) sorted from a bone marrow aspirate of a healthy donor. Active HSPCs (aHSPCs) were distinguished from quiescent HSPCs (qHSPCs) by presence/absence of Ki67 (filled and open arrowheads, respectively). Significant anticorrelation was observed between Ki67 mean intensity and nucleocytoplasmic ratio of poly(A) RNA (Pearson’s product moment correlation, correlation coefficient -0.19, p=0.0001; not shown). **b,** High magnification of aHSPC and qHSPC (arrowheads in low-magnification). Images are SUM intensity projections of Z-stack; scale bars: 10 µm. **c,** Quantifications from individual cells such as those depicted in a. Histograms represent mean and error bars s.e.m. of values normalised to aHSPC average. Mann-Whitney test, *p<0.05.

**Extended Data Fig 6.**
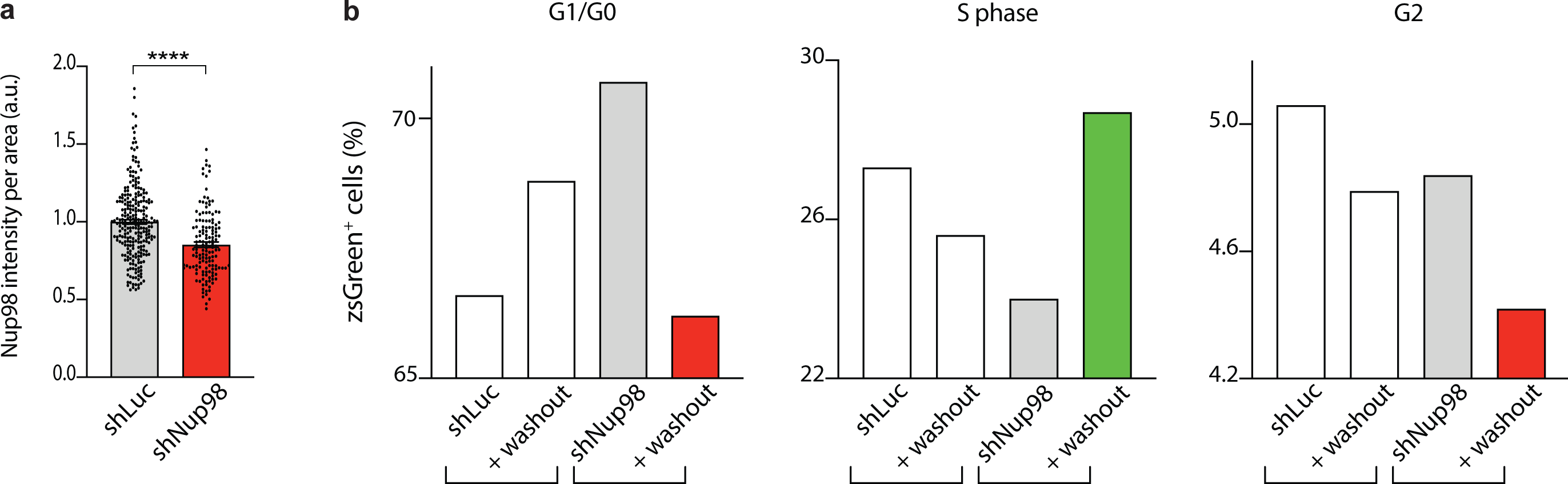
The effect of Nup98 downregulation on the cell cycle is reversible. **a**, Quantification of Nup98 levels in individual nuclei of cells expressing shRNAs after 3 d Doxycycline (Dox) induction. Histograms represent mean and error bars s.e.m. of values normalised to average of controls (shRNA targetting *Renilla* Luciferase, Luc). Mann-Whitney test, ****p<0.0001. **b**, Distribution of transduced cells (reported by zsGreen) according to DNA content as determined by flow cytometry after either 3 d Dox or 3 d Dox followed by 3 d washout.

**Extended Data Fig. 7.**
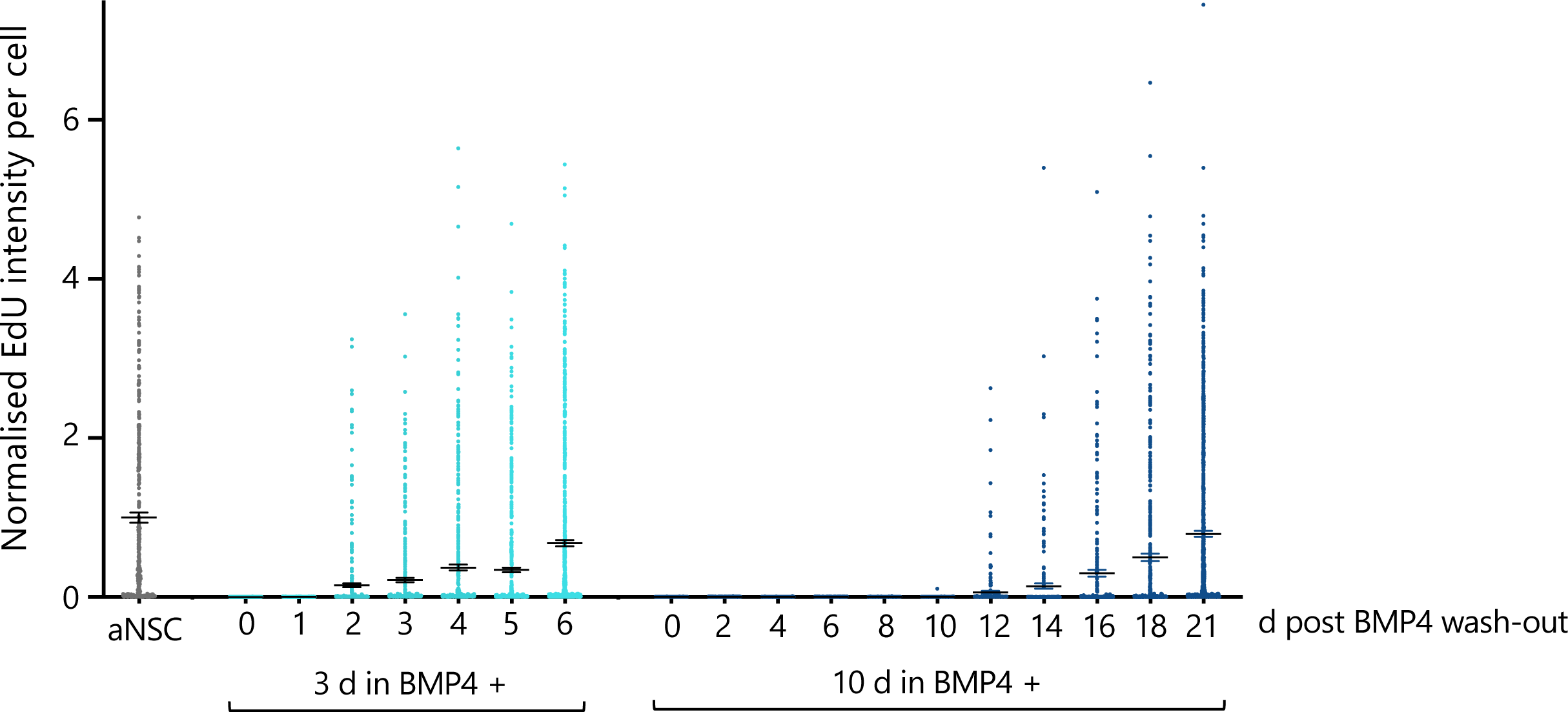
Longer exposure to BMP4 induces deeper quiescence in adult mouse hippocampal NSC cultures. EdU intensity was determined for individual nuclei and normalised to average of active condition. Following BMP4 wash-out, qNSCs that had been exposed to BMP4 for 3 d reactivated faster than those exposed to BMP4 for 10 d. Histograms represent mean and error bars s.e.m..

**Extended Data Fig. 8.**
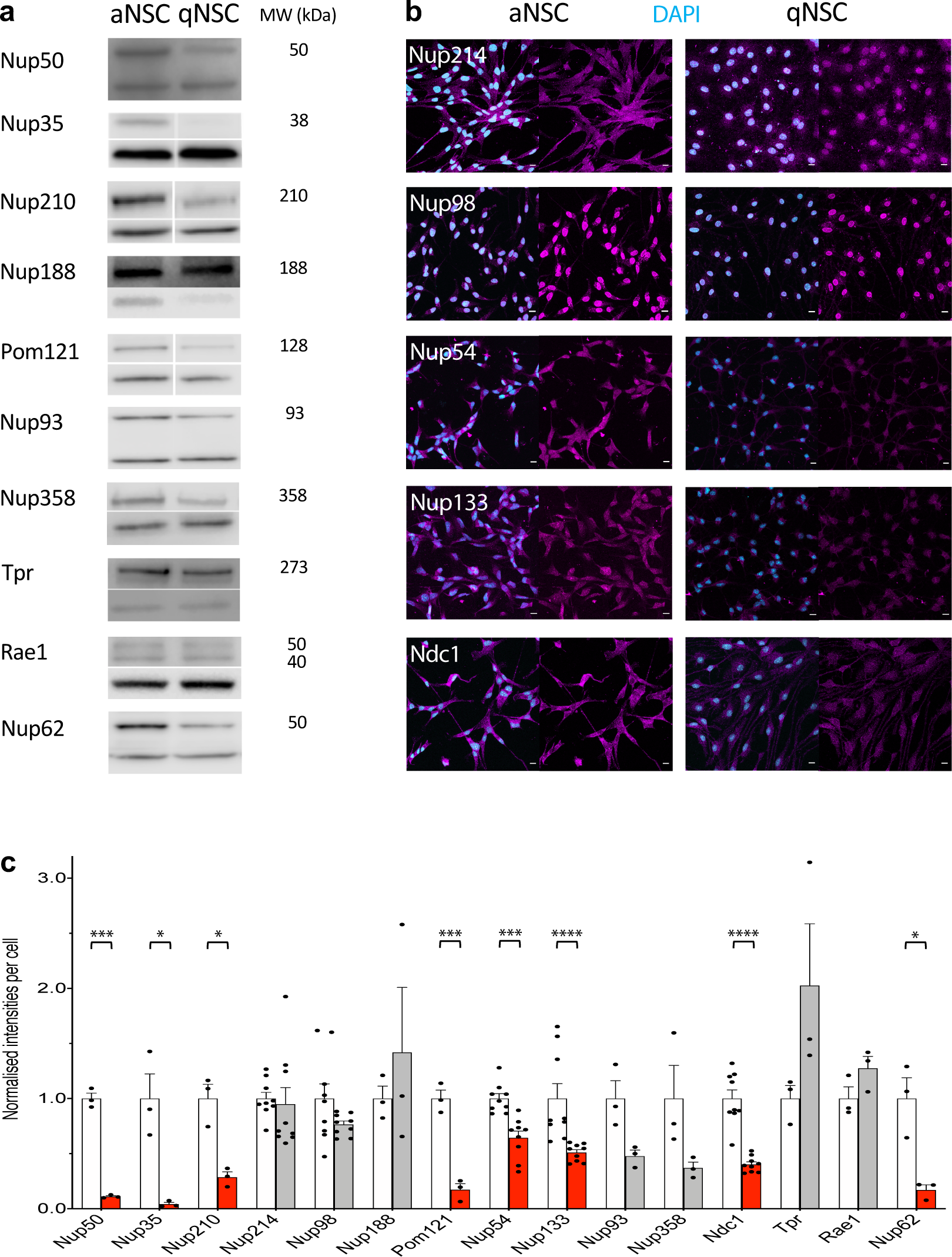
Assessment of nucleoporin downregulation in qNSCs relative to aNSCs. **a.** Nucleoporins assessed by Western blot (WB); for each, lower band shows Actin loading control at 42 kDa. Note that the observed molecular weight (MW) can vary from the predicted due to post translational modifications and experimental factors. **b**. Nucleoporins assessed by fluorescent immunocytochemistry (ICC). **c.**Quantification of samples such as those depicted in a-b ordered from most to least downregulated in proteomics (3 data points for WB, 9 for ICC). Histograms represent mean and error bars s.e.m. of values normalised to respective aNSC average (dark grey). Welch’s t test (WB): *p<0.05, **p<0.01, ***p<0.005; Mann-Whitney test (ICC): ***p<0.0005, ****p <0.0001.

**Extended Data Fig. 9.**
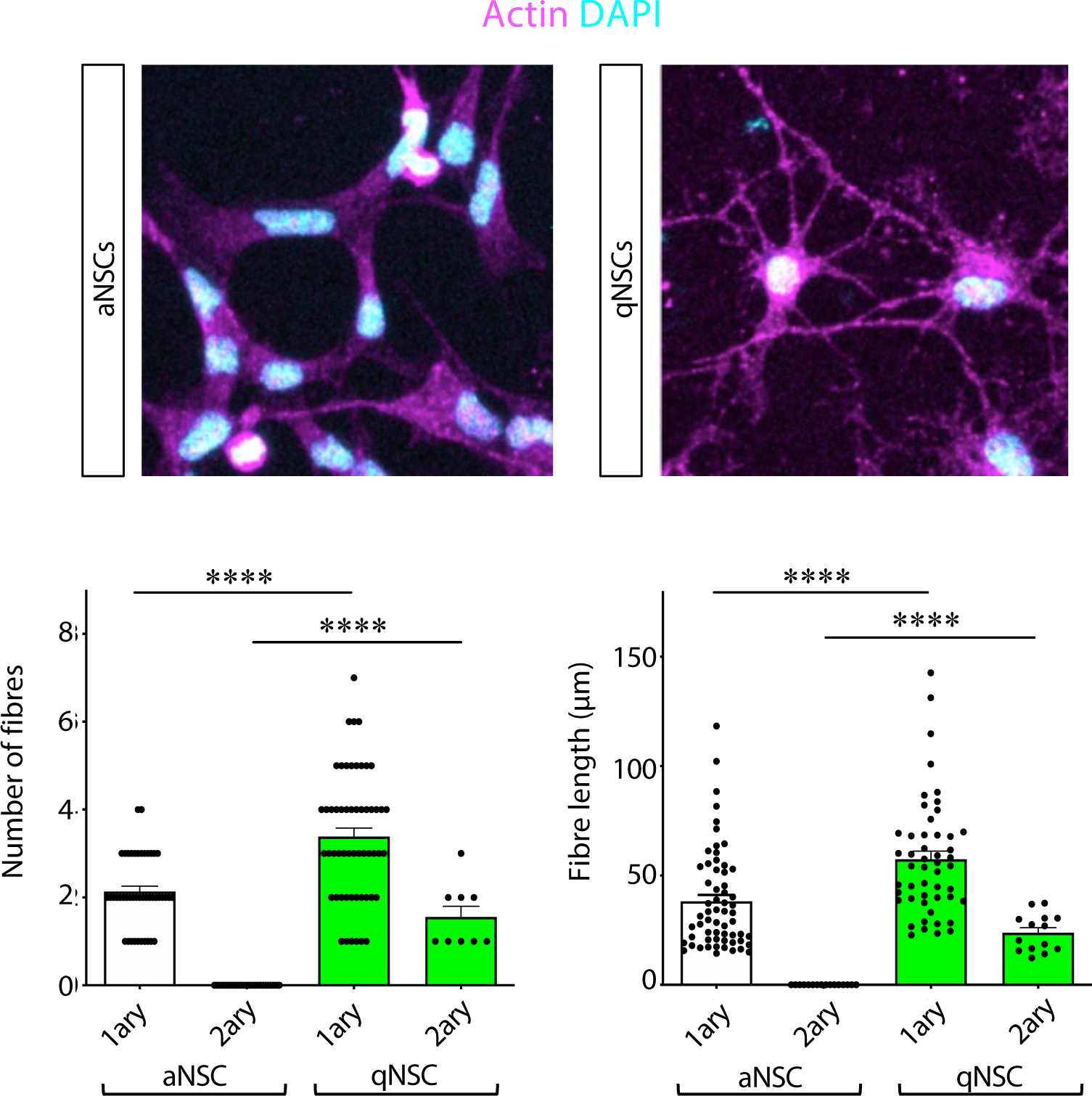
Mouse adult hippocampal NSCs in culture increase morphological complexity upon quiescence induction. Images are maximum intensity projections of a few optical sections. Histograms represent mean and error bars s.e.m.. Mann-Whitney test ****p<0.0001.

**Extended Data Fig. 10.**
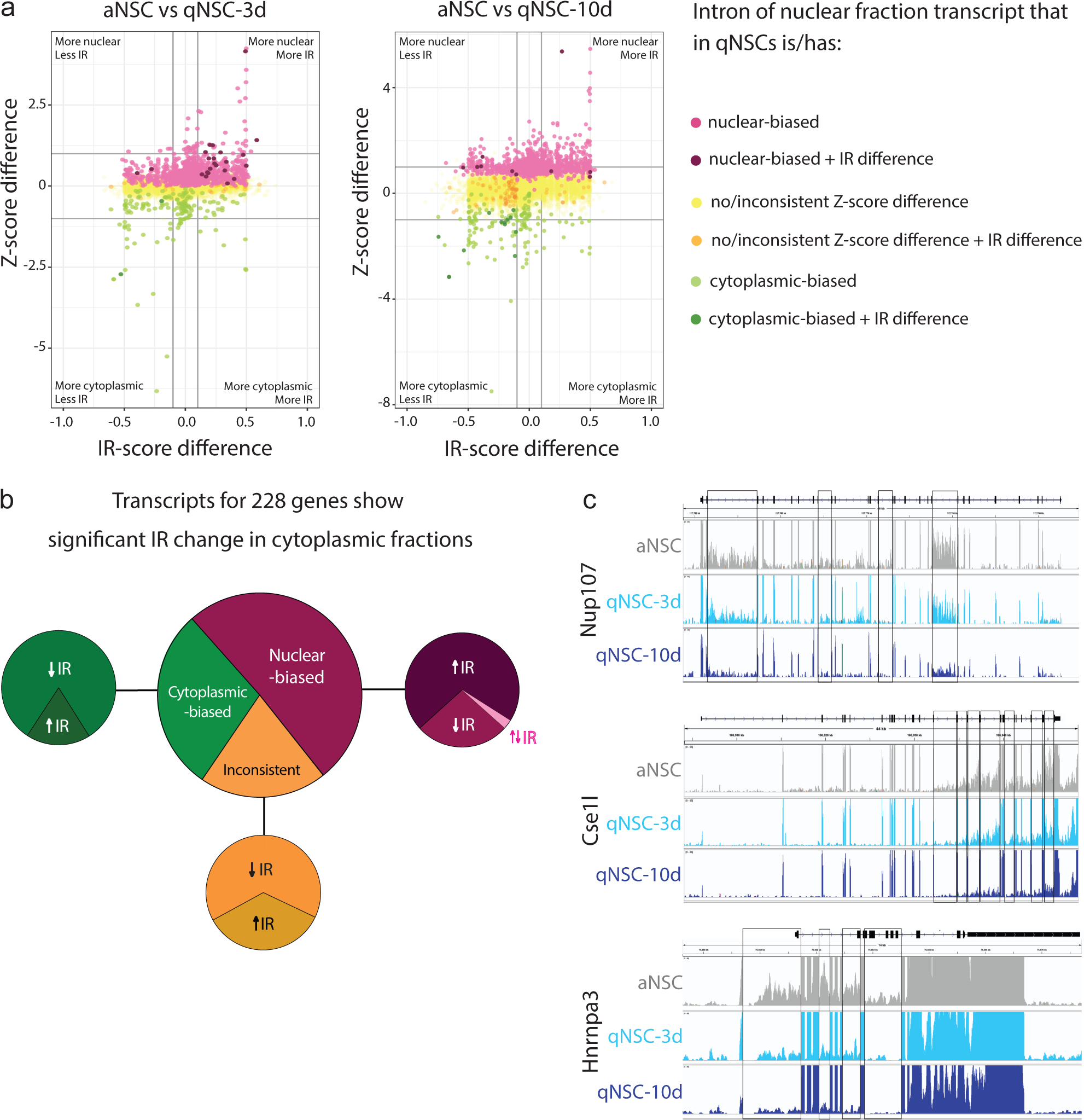
Nuclear-biased transcripts show increased intron retention in the cytoplasmic but not nuclear fraction of qNSCs. **a,b** depict data from cytoplasmic fractions. **a**. Plot of Z-score and IR-score differences between aNSCs and qNSCs per gene (single intron with largest IR difference plotted per gene). **b.** Large pie chart: breakdown of cytoplasmic fraction genes according to Z-score bias in qNSCs relative to aNSCs; small pie charts: breakdown of (un)biased genes according to direction of differential IR. **c**. The various introns of a transcript generally showed the same directionality of IR as NSCs shifted from active into deeper quiescence. Sequencing coverage tracks of three nuclear-biased transcripts in nuclear fractions for indicated conditions. Differential IR events (decreased in qNSCs) are highlighted by black boxes.

