## Supplemental Table 1 for "Neural stem cells alter nucleocytoplasmic partitioning and accumulate nuclear polyadenylated transcripts during quiescence"

**Supplemnetary Table 1 | Nucleoporin classification according to location on the NPC.** Grouped and colour-coded by subtype (Cautain B. *et al*. *FEBS Journal* 282, 445–462, 2015) as per the schematic. The novel *Drosophila* nucleoporin Snx was placed next to Nup98 given their sequence similarity but note that there is no evidence of what structural subtype it belongs to.

| Symmetric | | | | | |
|---|---|---|---|---|---|
| Scaffold Nups | | | | | |
| Coat Nups | | | Adaptor Nups | | |
| Human | *Drosophila* | Yeast | Human | *Drosophila* | Yeast |
| SEH1 | Nup44A | Seh1 | NUP93 | Nup93-1/2 | Nic96 |
| NUP75/85 | Nup75 | Nup85 | NUP205 | Nup205 | Nup192 |
| NUP160 | Nup160 | Nup120 | NUP188 | CG8771 | Nup188 |
| SEC13 | Sec13 | Sec13 | NUP155 | Nup154 | Nup157/170 |
| NUP96 | Nup96 | Nup145c | NUP35/53 | CG6540 | Nup53 |
| NUP107 | Nup107 | Nup84 | - | - | Nup59 |
| NUP133 | Nup133 | Nup133 | | | |
| NUP37 | Nup37 | - | | | |
| NUP43 | Nup43 | - | | | |

| | Asymmetric | | | | | | |
|---|---|---|---|---|---|---|---|
| | Cytoplasmic Nups | | | Nuclear basket Nups | | | |
| | Human | *Drosophila* | Yeast | Human | *Drosophila* | Yeast | |
| * | NUP98 | Nup98 | Nup116/100/145N | NUP98 | Nup98 | Nup116/100/145N | * |
| * | - | CG14712/Snx | - | | CG14712/Snx | - | * |
| * | NUP358 | Nup358 | - | NUP153 | Nup153 | Nup1 | * |
| * | NUP214 | Nup214 | Nup159 | NUP50 | Nup50 | Nup2 | * |
| * | NUPL2 | - | Nup42 | Tpr | Mtor | Mlp1/Mlp2 | |
| | NUP88 | Mbo | Nup82 | - | - | Nup60* | * |
| | GLE1 | CG14749 | Gle1 | - | - | Nup61 | |
| | RAE1 | Rae1 | Gle2 | | | | |
| | ALADIN | CG16892 | - | | | | |

| | Anchor / Poms / Transmembrane | | |
|---|---|---|---|
| | Human | *Drosophila* | Yeast |
| | NDC1 | Ndc1 | NDC1 |
| | Gp210 | Gp210 | - |
| * | POM121 | - | - |
| | - | - | POM34 |
| | - | - | POM152 |

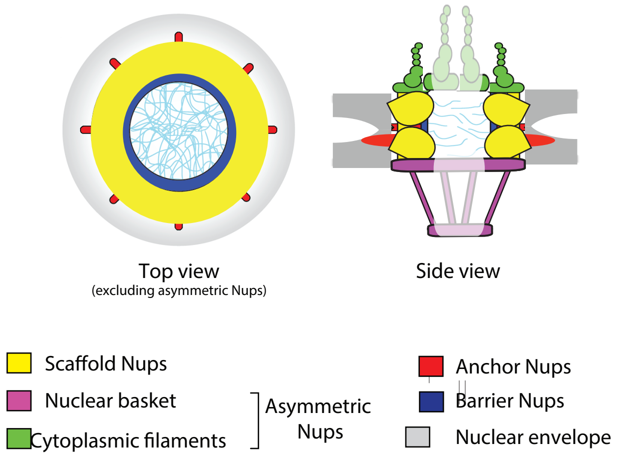


| | Mesh / Barrier FG Nups | | |
|---|---|---|---|
| | Human | *Drosophila* | Yeast |
| * | NUP62 | Nup62 | Nsp1 |
| * | NUP54 | Nup54 | Nup57 |
| * | NUPL1/NUP45 | Nup58 | Nup49 |

* **FG Nups: Phenylalanine (F) and Glycine (G) -rich**
