## Supplemental Table 2 for "Neural stem cells alter nucleocytoplasmic partitioning and accumulate nuclear polyadenylated transcripts during quiescence"

**Supplementary Table 2 | Karyopherin classification.** Karyopherins, also known as nuclear transport receptors or transportins, are grouped and colour-coded by directionality. Adapted from information contained in Kimura M. *et al. eLife* 6:e21184, 2017.

**Importins**

| | | Human | *Drosophila* |
|---|---|---|---|
| alpha | Importin-α1/2 | KPNA2 | Pen (Kap-α2) |
| | Importin-α3 | KPNA4 | Kap-α3 |
| | Importin-α4 | KPNA3 | αKap4 |
| | Importin-α5 | KPNA1 | Kap-α1 |
| | Importin-α6 | KPNA5 | " |
| | Importin-α7 | KNPA6 | " |
| beta | Importin-ß (Importin-ß1) | KNPB1 | Fs(2)Ket |
| | Importin-ß2 | TNPO1 (TRN-1) | Tnpo |
| | Transportin-2 | TNPO2 (TRN-2) | CG8219 |
| | Importin-12 | TNPO3 (TRN-SR, TRN-3) | Tnpo-SR |
| | Importin-4 | IPO4 (RANBP4) | CG32164/CG32165 |
| | Importin-5 (Importin-ß3) | IPO5 (RANBP5) | Karyß3 |
| | Importin-7 | IPO7 (RANBP7) | Msk |
| | Importin-8 | IPO8 (RANBP8) | " |
| | Importin-9 | IPO9 (RANBP9) | Ranbp9 |
| | Importin-11 | IPO11 (RANBP11) | Imp-ß11 |

**Exportins**

| | Human | *Drosophila* |
|---|---|---|
| RanBP3 | RANBP3 | RanBP3 |
| Exportin-1 | XPO1 (CRM1) | Emb |
| Exportin-2 | XPO2 (CAS, CSE1L) | Cse |
| Exportin-5 | XPO5 (RANBP21) | Ranbp21 |
| Exportin-6 | XPO6 (RANBP20) | Ebo |
| Exportin-7 | XPO7 (RANBP16) | Ranbp16 |
| Exportin-t | XPOT | - |
| RanBP17 | RANBP17 | - |

**Bidirectional**

| | Human | *Drosophila* |
|---|---|---|
| Exportin-4 | XPO4 | - |
| Importin-13 | IPO13 | Cdm |

**Undetermined directionality**

| | Human | *Drosophila* |
|---|---|---|
| RanBP6 | RANBP6 | - |
